# *Legionella* para-effectors target chromatin and promote bacterial replication

**DOI:** 10.1101/2022.10.16.512414

**Authors:** Daniel Schator, Sonia Mondino, Jérémy Berthelet, Cristina Di Silvestre, Mathilde Ben Assaya, Christophe Rusniok, Fernando Rodrigues-Lima, Annemarie Wehenkel, Carmen Buchrieser, Monica Rolando

## Abstract

*Legionella pneumophila* replicates intracellularly by secreting effectors *via* a type IV secretion system. One of these effectors is a eukaryotic methyltransferase (RomA) that methylates K14 of histone H3 (H3K14me3) to counteract host immune responses. However, it is not known how *L. pneumophila* infection catalyses H3K14 methylation as this residue is usually acetylated. Here we show that *L. pneumophila* secretes a eukaryotic-like histone deacetylase (LphD) that specifically targets H3K14ac and works in synergy with RomA. Both effectors target host chromatin and bind the HBO1 histone acetyltransferase complex that acetylates H3K14. Full activity of RomA is dependent on the presence of LphD as H3K14 methylation levels are significantly decreased in a Δ*lphD* mutant. The dependency of these two chromatin-modifying effectors on each other is further substantiated by mutational and virulence assays revealing that the presence of only one of these two effectors impairs intracellular replication, while a double knockout (Δ*lphDΔromA*) can restore intracellular replication. Uniquely, we present evidence for “para-effectors”, an effector pair, that actively and coordinately modify host histones to hijack the host response. The identification of epigenetic marks modulated by pathogens opens new vistas for the development of innovative therapeutic strategies to counteract bacterial infection and strengthening host defences.

## INTRODUCTION

The accessibility of chromatin to transcription factors and the subsequent changes in gene expression are key regulatory mechanism in eukaryotic cells. The process of changing this accessibility is known as chromatin remodeling. The dynamic modification of histones, the small basic proteins that DNA is wrapped around to form chromatin, is one of the most studied mechanisms of chromatin remodeling. The so-called histone tails – peptide sequences reaching out of the core histone structure – are subjected to a variety of post-translational modifications (PTMs) ^1^, such as acetylation, methylation, ubiquitination or phosphorylation ^2^. Methylation and acetylation of the amino-terminal tail of histone proteins are the most studied and best characterized ones. These modifications were the first histone PTMs discovered and were linked to altered rates of DNA transcription almost six decades ago ^3 4^. Importantly, combinations of acetylation and methylation of lysine residues in histone tails can function in a concerted manner with both cooperative and antagonistic functions. Methylation, depending on the target residue, can be associated to compaction of chromatin and reduced transcription ^5^ whereas acetylation often impairs the affinity of histones to DNA, consequently loosening chromatin compaction, promoting the recruitment of transcription factors ^6^ and increasing the mobility of histones along the DNA ^7^. Therefore, the balanced activity of enzyme classes involved in adding (histone methyltransferases and histone acetyltransferases (HAT)) and removing (histone demethylases and histone deacetylases (HDAC)) these groups ensure the correct expression of specific genes at specific times.

Numerous stimuli have been shown to influence the PTM levels of histones in the eukaryotic cell. Excitingly, it has been shown recently that different pathogens manipulate histone modifications mainly by secreting effector proteins targeting the nucleus, so-called nucleomodulins ^8^. In particular, histone methylation and histone acetylation have been shown to be potent targets for a variety of pathogens to promote their replication in their host cells ^9 10 11 12^. One of the pathogens described to modulate histone PTMs of its host cell is *Legionella pneumophila,* a facultative intracellular, Gram-negative bacterium that parasitizes free-living protozoa, but that is also the causative agent of a severe atypical pneumonia in humans, called Legionnaires’ disease ^13^. The intimate interaction of *L.pneumophila* with its eukaryotic hosts has shaped the bacterial genome and led to the evolution of numerous mechanisms allowing *L. pneumophila* to manipulate host functions and to thrive in this otherwise hostile intracellular environment. This *Legionella*-protozoa-coevolution is particularly reflected in the presence of multiple genes encoding eukaryotic-like proteins in the *Legionella* genome ^14 15 16^. Many of these proteins are secreted by the Dot/Icm type-IV secretion system (T4SS) that translocates more than 300 proteins into the host cell that are key for facilitating bacterial intracellular survival and replication ^17 18 16 19^. One of these secreted eukaryotic-like effectors is a SET-domain histone methyltransferase, RomA, that mimics eukaryotic histone methyltransferases and directly modifies host chromatin by tri-methylating lysine 14 on histone H3 (H3K14) ^20^. The unique activity of this bacterial protein leads to an increase in H3K14 methylation thereby decreasing the expression of many genes involved in the host response to the infection ^20^. Interestingly, H3K14me had not been identified in human cells before RomA activity was discovered but this modification was recently reported to be present at very low levels in human cells, probably explaining why it had been overlooked ^21^. H3K14me increases under specific conditions such as stress response ^22^; however, the prevalent modification of H3K14 is acetylation.

Thus, the question emerges “How does the bacterial pathogen *L.pneumophila* methylate a histone residue that is usually acetylated?”. Given the plethora of eukaryotic-like proteins in *L. pneumophila* we hypothesized that the bacteria might encode its own eukaryotic-like histone deacetylase (HDAC). Here we report that *L. pneumophila* encodes an HDAC-like protein (LphD) that works in synergy with RomA by specifically deacetylating H3K14 to facilitate methylation of H3K14 by RomA. Together these effectors function as effector pairs, or ‘para-effectors’, with high levels of interdependence that serve to fine tune the host cell’s gene expression and promote bacterial intracellular replication.

## RESULTS

### *L. pneumophila* encodes an HDAC-like protein that preferentially targets H3K14

Bioinformatic analyses of the *L. pneumophila* strain Paris genome identified the gene *lpp2163* encoding a 424 amino acid long protein, predicted to be a Zn^2+^-dependent histone deacetylase, that we named LphD (*<u>L</u>egionella <u>p</u>neumophila* <u>h</u>istone <u>D</u>eacetylase) ^12^. We computed a structural model of LphD using AlphaFold ^23^ which allowed prediction of the protein structure with high confidence (**Figure 1A**) except for the N- and C-terminal regions (amino acids 1-25 and 415-424), that are likely unstructured in solution. We sequence aligned a selection of eukaryotic HDACs (**Figure S1A**) to compute the conservation scores using ConSurf ^24^ and map them onto the LphD model **(Figure 1B)**. All the catalytic residues of the so-called charge relay system ^25^ including the active-site tyrosine (Y392) and the Zn^2+^ coordinating atoms were conserved (**Figure 1B inset**). An exception is N219 which is a histidine in eukaryotic structures but fulfills the same function in coordinating the zinc atom.

**Figure 1:**
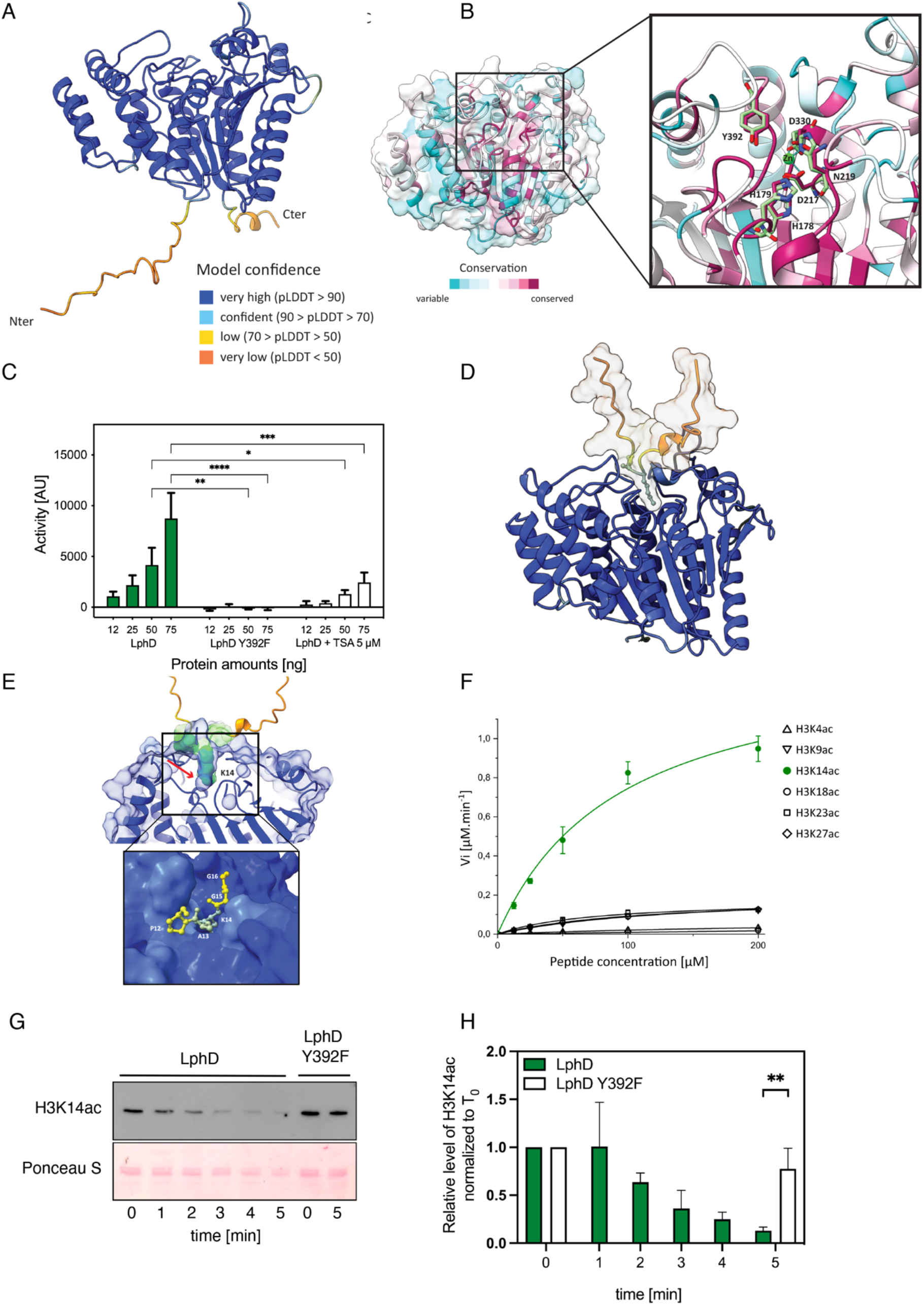
LphD has predicted structural similarity to eukaryotic HDACs and possesses deacetylase activity. (**A**) Cartoon representation of the LphD model generated by AlphaFold. The per-residue confidence score (pLDDT) produced by AlphaFold is shown in the insert. (**B**) The conservation score between LphD and representatives of the eukaryotic HDAC families is mapped onto the LphD model. The insert shows the substrate binding groove with the catalytic residues (binding pocket residues and catalytic center) of LphD highlighted and colored according to conservation and HDAC6 colored green (PDB 5EDU). The catalytic zinc is shown from the HDAC6 structure. (**C**) Fluor de Lys^®^HDAC activity assay of LphD. Increasing amounts of purified His-LphD and its predicted catalytic inactive mutant (His-LphD Y392F) were used to assess the lysine deacetylase activity *in vitro*. 5μM of Trichostatin A (TSA), a broad range HDAC inhibitor, was added to LphD as a control (n = 3 ± SD, * p < 0.05, ** p < 0.01, *** p < 0.001, **** p < 0.0001). (**D**) Cartoon representation of the LphD-H3 peptide complex generated by AlphaFold. The H3 peptide is represented as a surface/cartoon model, with the K14, A15 and P16 residues shown as sticks. (**E**) The H3K14 (in green) is positioned towards the active site and the active site cavity (in blue) shows space to accommodate an acetylated K14 residue (red arrow). **Inset**: Residues 12 to 16 of the H3 peptide are represented as ball and stick, the entrance of the active site of LphD is represented as surface. (**F**) Steady-state kinetics of purified LphD on fluorescent H3 peptide substrates acetylated at different lysine residues (H3K4ac, H3K9ac, H3K14ac, H3K18ac, H3K23ac, H3K27ac) (n = 3 ± SD). V_i_values (μM.min^-1^) were plotted against substrate peptide concentrations and curves were fitted using Michaelis–Menten equation. (**G**) Representative western blot of LphD activity on H3K14ac levels on high-acetylated histone octamers. Wild type protein (LphD) is compared to the catalytic inactive mutant (LphD Y392F). H3K14ac levels are assessed every minute for 5 minutes by immunoblot. Ponceau S staining of H3 is used as loading control. (**H**) Densiometry quantification of LphD activity on H3K14ac levels on high-acetylated histone octamers (same conditions as in **G**). Octamers are incubated with LphD (green) and LphD Y392F (white) and the reaction was stopped after the indicated time (in minutes). The H3K14ac signal was quantified after immunoblot detection and normalized to the signal at 0 min (n = 3 ± SD, ** p < 0.01).

To determine if LphD indeed possesses lysine deacetylase activity, we performed a fluorometric enzymatic assay that allows the quantification of lysine deacetylation on an acetylated lysin side chain. **Figure 1C** shows that LphD exhibits *in vitro* lysine deacetylase activity in a dose-dependent manner. Furthermore, the single amino acid substitution (Y392F) at the predicted active site completely inactivates the enzyme. Adding Trichostatin A (TSA), a broad range inhibitor of Zn^2+^-dependent histone deacetylases, reduced LphD activity (**Figure 1C**), further suggesting that LphD is a Zn^2+^-dependent histone deacetylase. We then attempted to identify the targeted lysine(s) by using MS/MS analyses of histones extracted from THP-1 cells that had been incubated with purified LphD for 1h at 37°C. Several lysine residues were identified on H3, H4 and H2B (**Figure S1B**). However, as this approach did not use physiological quantities of LphD, it might not have revealed the specificity this protein could have during infection. Thus, we used AlphaFold to compute models of the complexes between LphD and the tails of histones H3 and H4 (residues 1-25) to see if we could get an indication for the physiological substrate specificity. In the case of H3, all 5 models show the same lysine (H3K14) binding to this pocket (**Figure S1C**). In contrast, for the H4 peptide the models were predicted with poor confidence and in only 2 out of 5 complexes is a lysine (H4K9) residue positioned at the active site (**Figure S1D**). Furthermore, whereas most of the H3 peptide is predicted with poor confidence and away from the LphD surface, H3K14 is placed into the active site pocket, a tight cavity that accommodates H3K14 and shows room for an additional acetyl group (**Figure 1D** and **1E**). None of the other lysines of the peptide were predicted to bind LphD, probably due to specific substrate recognition through the flanking amino acids Pro12, Ala13 and Gly15, Gly16 (**Figure 1E inset**). Based on the results of these models, H3K14 seems to be the preferred target of LphD.

We first analysed *in vitro* if LphD has indeed target specificity for H3K14 as predicted. Using histone H3 peptides acetylated on each lysine residues present on the H3 tail (H3K4, H3K9, H3K14, H3K18, H3K23 and H3K27) we measured the catalytic efficiency of LphD with varying peptide concentrations and with LphD-Y392F as control. As seen in **Figure 1F** LphD has a high catalytic efficiency for H3K14 with an efficiency constant (k_cat_/K_M_) of 1.984 μM^-1^ .min^-1^ as compared to a k_cat_/K_M_ of below 0.314 μM^-1^ .min^-1^ for all other residues (**Table 1**). Deacetylase activity of LphD on H3K14 was also observed when testing histone octamers isolated from human cells (**Figure 1G**, **1H**) with H3K14ac antibodies that we had validated by dot blot (**Figure S1E**). Indeed, the value of kcat/KM. is the highest for H3K14 and the deacetylation of H3K14 on octamers occurred already within 5 min, whereas for longer incubation times a loss of specificity was observed. This explains the MS/MS results obtained previously, where a one-hour incubation time led to a deacetylation of many residues and a loss of specificity.

**Table 1:** Enzymatic constants of LphD activity against different acetylated peptides

| Substrate | $K_M [\mu M]$ | $k_{cat} [min^{-1}]$ | $k_{cat}/K_M [\mu M^{-1}.min^{-1}]$ |
| --- | --- | --- | --- |
| H3K4ac | 286.96 | 10.34 | 0.036 |
| H3K9ac | 143.15 | 28.06 | 0.196 |
| H3K14ac | 95.13 | 188.78 | 1.984 |
| H3K18ac | 3037.16 | 36.20 | 0.011 |
| H3K23ac | 73.74 | 23.22 | 0.314 |
| H3K27ac | 114.08 | 25.58 | 0.224 |

Kinetics constants ( $K_M$ and $k_{cat}$ ) were individually calculated for each peptide from non-linear regression fit using OriginPro.

Taken together, AlphaFold models, enzymatic assays on acetylated histone peptides as well as the use of specific antibodies revealed that LphD has clear target specificity for H3K14.

### LphD is a secreted effector targeting host histones during infection

To characterize the function of LphD during infection we first assessed whether it is a T4SS substrate. THP-1 cells were infected with an *L. pneumophila* strain expressing a β-lactamase-LphD fusion construct or the-β-lactamase-RomA (positive control) and translocation was measured by quantifying the number of cells exhibiting β-lactamase activity against a fluorescent substrate (CCF4). The secretion of β-lactamase-LphD was clearly detected during infection. Moreover, LphD is secreted by the T4SS, as no β-lactamase activity was observed when the β-lactamase-LphD fusion protein was produced by a strain lacking a functional Dot/Icm secretion system (Δ*dotA*) (**Figure 2A and S2A**). To assess the subcellular localization of LphD, we transiently transfected HeLa cells with an EGFP-LphD fusion product, showing that it accumulates in the nucleus of transfected cells (**Figure 2B**), compared to the cytosolic localization typically found for EGFP (**Figure S2B**). Furthermore, in LphD transfected cells a drastic decrease of the H3K14ac signal occurred, which was not seen in cells transfected with the Y392F mutant, although the LphD-Y392F catalytic inactive mutant was also located in the nucleus of transfected cells (**Figure 2C**). An antibody raised against LphD (**Figure S2C**) confirmed that LphD accumulates in the host cell nucleus also during infection (**Figure 2D**).

**Figure 2:**
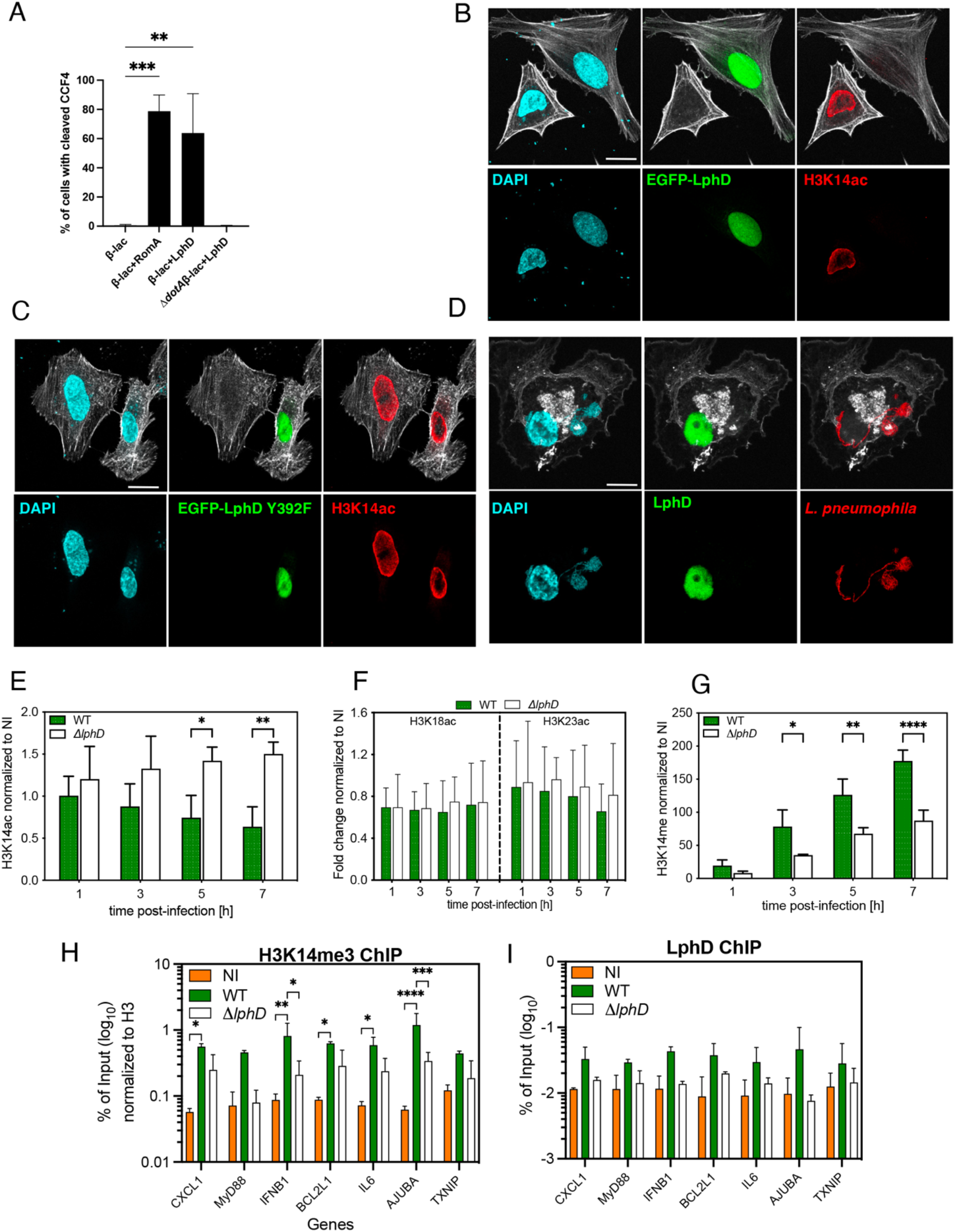
LphD is a secreted effector that targets H3K14 in the nucleus. (**A**) β-lactamase secretion assay for LphD. Percentage of cells with cleaved CCF4 were calculated as a ratio of secretion-positive blue cells over total stained cells (green and blue positive) after 2 hours of infection with *L.pneumophila* wild type or Δ*dotA* overexpressing expressing either β-lactamase alone (β-lac), β-lactamase fused to LphD (β-lac+LphD) or β-lactamase fused to RomA (β-lac+RomA) (n = 3, ± SD, ** p < 0.01; *** p < 0.001). (**B-C**) Immunofluorescence analysis of subcellular localization of EGFP-LphD and H3K14 acetylation. HeLa cells were either transfected with EGFP-LphD (B) or EGFP-LphD Y392F (C) for 24hours and then stained for H3K14ac using a specific antibody. DAPI (cyan), EGFP-LphD/LphD Y392F (green), H3K14ac (red), and phalloidin (gray). Single channel images are shown. Scale bars 10 μm. (**D**) Immunofluorescence analysis of subcellular localization of LphD during infection. Differentiated THP-1 cells were infected 16 hours at an MOI of 50 with *L. pneumophila* wild type expressing LphD and DsRed. DAPI (cyan), LphD (green), *L. pneumophila* (red), and phalloidin (gray). Scale bars 10 μm. (**E** and **F**) Quantification of western blot signal of acetylation of H3K14 (**E**) and H3K18 and H3K23 (**F**). THP-1 cells were infected with *L. pneumophila* wild type (green) or Δ*lphD* (white) strains expressing EGFP. Cells were sorted by FACS based on the EGFP signal at different times post-infection, as indicated, to enrich for infected cells. Histones were isolated by acidic extraction and analysed by western blot. Histone H1 was used as loading control and signal is fold change normalized to non-infected cells (NI) (n = 3 ± SD, * p < 0.05, ** p < 0.01). (**G**) Quantification of western blot signals of H3K14 methylation. THP-1 cells were infected with *L. pneumophila* wild type (green) or Δ*lphD* (white) strain expressing EGFP. Cells were sorted by FACS based on the EGFP signal at different times post-infection, as indicated, in order to enrich for infected cells. Histones were isolated by acidic extraction and analysed by western blot. Histone H1 was used as loading control and signal is compared to non-infected cells (n = 3 ± SD, * p < 0.05, ** p < 0.01, **** p< 0.0001). (**H**) Chromatin immunoprecipitation of H3K14me during *L.pneumophila* infection followed by qPCR targeting selected promoters. THP-1 cells were infected 7 hours with *L. pneumophila* wild type (green) or Δ*lphD* (white) strains expressing EGFP. Cells were sorted based on EGFP signal by FACS in order to enrich for infected cells. Chromatin was isolated and H3K14me IP was performed. Signal was normalized to histone H3 and compared to non-infected cells (NI, orange) (n = 2 ± SD, * p < 0.05, ** p < 0.01, *** p<0.001, **** p < 0.0001). (**I**) Chromatin immunoprecipitation of LphD during *L. pneumophila* infection followed by qPCR targeting selected promoters. THP-1 cells were infected with *L. pneumophila* wild type (green) or Δ*lphD* (white) strains expressing EGFP. Cells were sorted at different times post-infection by FACS. Chromatin was isolated and LphD IP was performed. Signal compared to non-infected cells (NI, orange) (n = 2 ± SD).

To assess the influence of LphD on the epigenetic status of H3K14 during infection, we isolated histones from cells infected with either *L. pneumophila* wild type or a Δ*lphD* strain and followed H3K14 acetylation as a function of time. **Figure 2E** shows that *L. pneumophila* wild type infection leads to a decrease in H3K14ac within 7 hours of infection, dependent on the presence of LphD, as the infection with the Δ*lphD* strain led to an increase in H3K14ac in the same timeframe (**Figure 2E and S2D**). LphD activity during infection is specific for H3K14, as other tested residues (H3K18 and H3K23) did not show a difference whether the cells had been infected with the wild type or the Δ*lphD* mutant strain (**Figure 2F, S2E and S2F)**. Thus *L. pneumophila* encodes two effectors targeting the same lysine on the same histone tail, RomA that targets and methylates H3K14 ^20^ and LphD that deacetylates H3K14. As we observed a decrease in H3K14 acetylation in presence of RomA, we wondered whether this was due to a genome wide H3K14me accumulation, or a specific and targeted deacetylase activity, possibly driven by the bacteria. Indeed, when LphD is absent (Δ*lphD*) the level of H3K14me during infection is significantly reduced from the early stages of infection (1-3 hours) (**Figure 2G and S2G**). To analyse whether LphD and RomA can act at the same promoters, we analysed promoters of genes we had previously shown to be the targets of H3K14 methylation by RomA by ChIP analyses ^20^. This revealed that the methylation levels of H3K14 on specific promoters significantly decrease when infecting with a Δ*lphD* mutant strain where LphD deacetylation activity is missing, compared to wild type infection (**Figure 2H**). Thus, methylation of H3K14 at these promoters is due to RomA activity, but it also needs LphD binding to the same regions of the chromatin (**Figure 2I**).

These results suggests that LphD directly impacts the activity of RomA on histones, and strongly indicates that the two effectors act in concert to modify the host chromatin landscape and downregulate host defense gene expression.

### LphD and RomA have complementary functions as virulence factors

Replication of the wild type strain compared to the Δ*lphD* mutant strain in macrophages derived from THP-1 cells as well as in *Acanthamoeba castellanii,* a natural host of *L. pneumophila,* was compared to determine whether LphD was important for *L. pneumophila* intracellular growth. The Δ*lphD* strain has a consistent growth defect compared to the wild type strain in both THP-1 cells and in *A. castellanii* (**Figure 3A** and **3B**). Complementation of the Δ*lphD* strain with full length *lphD* under the control of its native promotor completely reversed the growth defect and even induced a slight increase in replication due to the plasmid copy number, further underlining the role of LphD in virulence of *L. pneumophila* **(Figure 3C)**. When either LphD or RomA is deleted (Δ*lphD* and Δ*romA*) the bacteria show a defect in intracellular replication in THP-1 cells **(Figure 3D)** and in *A. castallanii* **(Figure 3E)**. However when the infection is performed with the double mutant Δ*lphD ΔromA* the phenotype is partially reversed **(Figure 3D and 3E)**, suggesting that the replication defect of the single knockout strains is not only caused by the absence of one effector, but also by the activity of the other effector alone. Hence, the advantageous influence of each effector on intracellular replication depends on the presence of the other. Moreover, this collaborative effect is dependent on the catalytic activity of each effector. Indeed, complementation of the double mutant with either *lphD* or *romA* alone shows the growth phenotype of the single mutant, which is not the case when complementing it with the catalytically inactive proteins (**Figure 3F**). This dependence is also seen when the transcriptional response of THP-1 cells infected with *L. pneumophila* wild-type is compared to that of THP-1 cells infected with the Δ*lphD, ΔromA* and Δ*lphDΔromA* knockout strains by RNAseq. In contrast to the Δ*romA* mutant, the transcriptional profile of the double knockout strain strongly resembles wild type infected cells (**Figure S3A)**.

**Figure 3:**
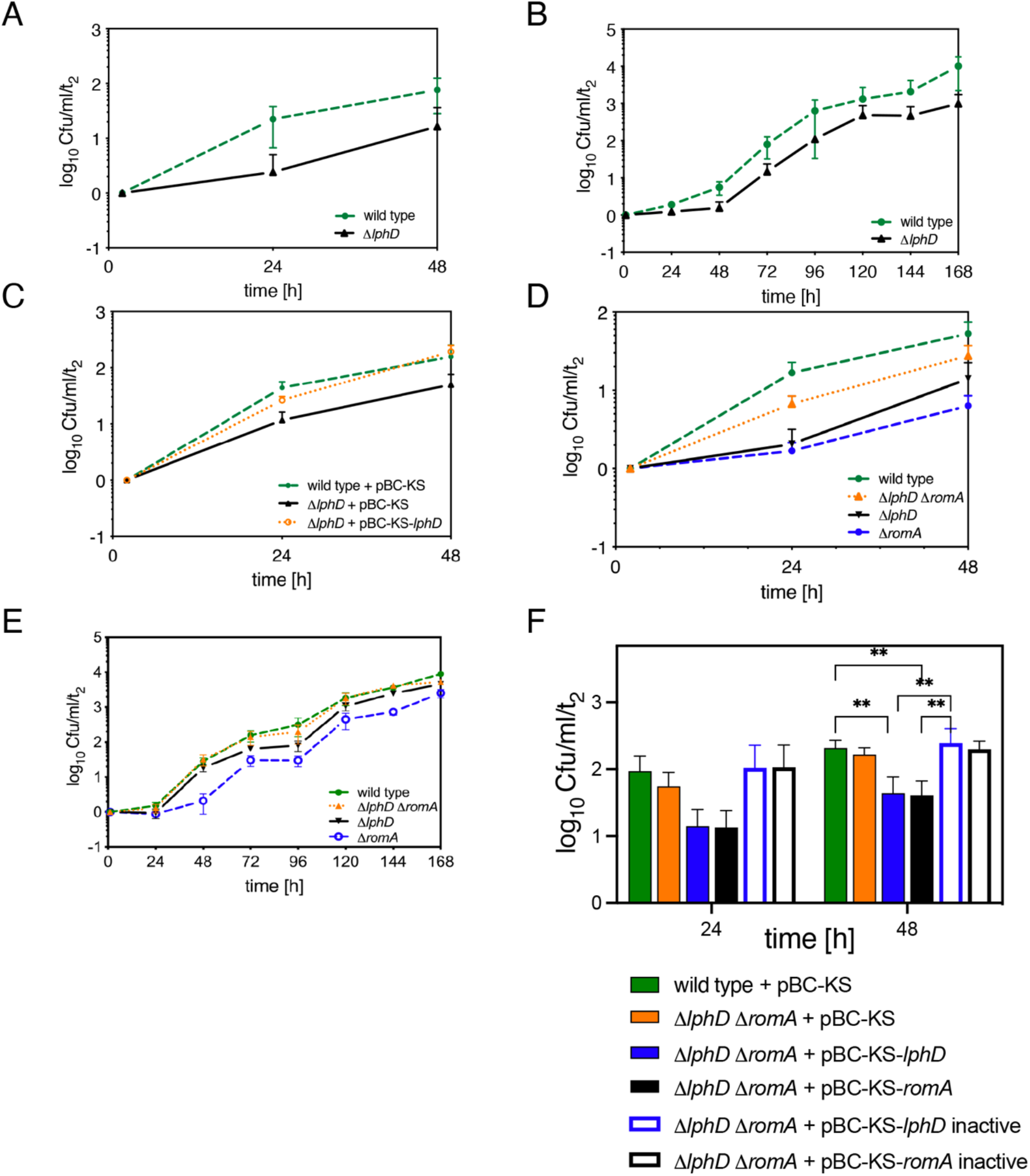
LphD and RomA act as para-effectors during intracellular replication. (**A**) Intracellular replication in THP-1 cells (MOI = 10) of wild type *L. pneumophila* (dashed green line) and the Δ*lphD* mutant strain (solid black line). Replication was determined by recording the number of colony-forming units (cfu) through plating on buffered charcoal yeast extract (BCYE) agar (log_10_ ratio cfu/ml/t_2_, n = 3 ± SD). (**B**) Intracellular replication in *A. castellanii* (MOI = 0.1) of wild type *L. pneumophila* (dashed green line) and Δ*lphD* mutant strain (solid black line). Replication was determined by recording the number of colony-forming units (cfu) through plating on buffered charcoal yeast extract (BCYE) agar (log_10_ ratio cfu/ml/t_2_, n = 3 ± SD). (**C)**Complementation analysis after infection of THP-1 cells (MOI = 10) with the *L. pneumophila* wild type and Δ*lphD* mutant carrying the empty vector pBC-KS (dashed green and solid black lines, respectively), or Δ*lphD* carrying the complementing plasmid pBC-KS-*lphD* (dotted orange) (log_10_ ratio cfu/ml/t_2_, n = 4 ± SD). (**D**) Intracellular replication in THP-1 cells (MOI = 10) of wild type *L. pneumophila* (dashed green line), the Δ*lphDΔromA* (orange dotted line), the Δ*lphD* (solid black line), and the Δ*romA* (dashed blue line) mutant. Replication was determined by recording the number of colony-forming units (cfu) through plating on buffered charcoal yeast extract (BCYE) agar (log_10_ ratio cfu/ml/t_2_, n = 4 ± SD) (**E**) Intracellular replication in *A. castellanii*(MOI = 0.1) of wild type *L. pneumophila*(dashed green line), Δ*lphD*Δ*romA* (dotted orange line,), Δ*lphD* (solid black line), and Δ*romA* (dashed blue line) mutant. Replication was determined by recording the number of colony-forming units (cfu) through plating on buffered charcoal yeast extract (BCYE) agar (log_10_ ratio cfu/t_2_, n = 3 ± SD.). (**F**) Bar plot of the complementation analysis after infection of THP-1 cells (MOI = 10) with the *L. pneumophila* wild type and Δ*lphD*Δ*romA* mutant carrying the empty vector pBC-KS (green and orange, respectively), the complementing plasmid pBC-KS-*lphD* (blue) or pBC-KS-*romA* (black), as well as their catalytic inactive counterparts pBC-KS-*lphD* Y392F (white, blue border) and pBC-KS-*romA* Y249F/R207G/N210A (white, black border) (log_10_ ratio cfu/ml/t_2_, n = 2 ± SD ** p < 0.01).

Taken together, RomA activity strongly depends on the presence of active LphD on the transcriptional level as well as in intracellular replication. Given the strong interdependence of these two effectors we coined the term “para-effectors” for the RomA and LphD pair.

### LphD and RomA target host chromatin cooperatively

As LphD and RomA target the chromatin together, we first tested if the two effector proteins interact. *In vitro* binding assays with His- and GST-tagged proteins produced in *Escherichia coli* showed that RomA and LphD can indeed directly interact (**Figure 4A)**. Furthermore, when expressed in eukaryotic cells, RomA and both LphD and LphD Y392F interact and immunoprecipitate with histone H3 proteins (**Figure 4B,** for IP control see **Figure S3B)**, confirming that RomA and LphD target the chromatin together. We thus searched for possible eukaryotic interacting partners or complexes that may facilitate chromatin targeting by these two bacterial effectors. To identify potential targets of LphD, we performed affinity chromatography by GFP-trap pull down of HEK293T cells transfected with an EGFP-LphD construct followed by MS/MS analysis. We set a threshold for candidate proteins that at least two unique peptides were detected with a significant false discovery rate (<0.1) and a high (>4) fold change as compared to the control condition (GFP). This approach identified a total of 542 significantly enriched proteins, compared to the control (EGFP) (**Figure 4C**). Interestingly, among this set of proteins, many of the identified peptides were derived from proteins that are known to be involved in epigenetic regulations of the cell (**Table S1**). Among those the histone acetyl transferase (HAT) KAT7 (HBO1) was a promising candidate; KAT7 is the enzymatic subunit of the so-called HBO1 complex, comprised of KAT7, BRPF1-3, ING4/5 and MEAF6. This complex is well-known to bind histone H3, regulating the acetylation of K14 (**Figure 4D**) ^26^.

**Figure 4:**
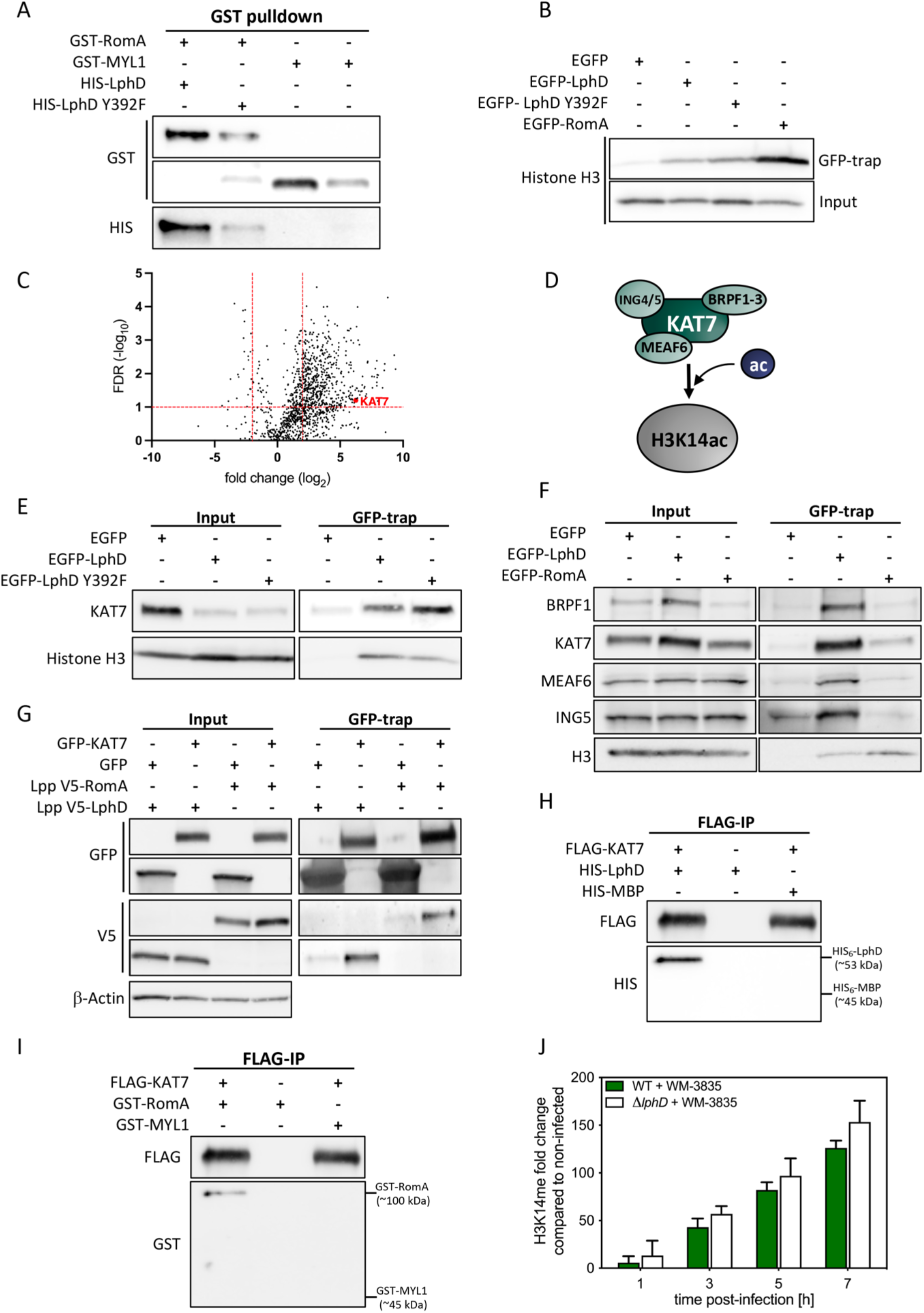
LphD and RomA taget chromatin *via* the HBO1/KAT7 complex. (**A**) *In-vitro* protein interaction assay using purified GST-RomA and HIS_6_-LphD. Purified proteins were mixed in equal amounts followed by a GST-pulldown using Glutathione-beads. GST-MYL1 was used as negative binding controls. (**B**) Immunoblots showing the interaction of LphD and RomA with histone H3. GFP-trap beads in HEK293T cells transfected with EGFP, EGFP-LphD, EGFP-LphD Y392F or EGFP-RomA. Pulldown samples were analyzed for the presence of H3. Input shows the expression level of endogenous histone H3 total lysates. For IP control see Figure S3B. (**C**) Volcano plot of EGFP-LphD interacting proteins. The log_2_ fold change of EGFP-LphD to control (GFP) is plotted against the −log_10_ of the false discovery rate (FDR). Protein selected for further tests (KAT7) is highlighted in red. Thresholds were set at log_2_ fold change > 2 and false discovery rate <0.1 (dashed red lines). (**D**) Schematic representation of the HBO1 histone acetyltransferase complex in the configuration with BRPF1-3 targeting H3K14. (**E**) Immunoblots showing the interaction of LphD with KAT7 and histone H3. GFP-trap beads (GFP-trap) in HEK293T cells transfected with EGFP, EGFP-LphD, or EGFP-LphD Y392F. Input shows the expression level of endogenous KAT7 and Histone H3 in total lysates. For IP control see Figure S3C. (**F**) Immunoblots showing the interaction of LphD and RomA with the HBO1 complex and histone H3. GFP-trap beads in HEK293T cells transfected with EGFP, EGFP-LphD, or EGFP-RomA. Samples were analyzed for the presence of the different components of the HBO1 complex (BRPF1, KAT7, MEAF6, ING5). Input shows the expression level of endogenous HBO1 components and histone H3 in total lysates. For IP control see Figure S3E. (**G**) Immunoblots showing the interaction of LphD and RomA with KAT7. GFP-trap beads in HEK293T cells transfected with either EGFP or EGFP-KAT7 followed by infection with *L. pneumophila* wild type over-expressing either V5-LphD or V5-RomA. EGFP proteins were pulled down using GFP-trap beads and samples were analyzed for the presence of V5-LphD or V5-RomA. Input shows the expression level in total lysates of transfected EGFP-KAT7, V5-fusion proteins, as well as β-actin (loading control). (**H** and **I**) *In-vitro* protein interaction assay using purified FLAG-KAT7, HIS_6_-LphD (**H**) and GST-RomA (**I**). Purified proteins were mixed in equal amounts followed by a FLAG-pulldown using FLAG-trap beads. HIS6-MBP and GST-MYL1 were used as negative binding controls. (**J**) Quantification of western blot signal for H3K14 methylation in WM-3835 (KAT7-specific inhibitor) treated cells. THP-1 cells were treated with WM-3835 (1 μM) 18 hours prior to infection with *L.pneumophila* wild type (green) or Δ*lphD* (white) strain expressing EGFP. Cells were sorted at different times post-infection by FACS, histones were isolated and analysed by western blot. Histone H1 was used as loading control and signal is compared to non-infected cells (n = 2 ± SD).

Using a GFP-trap to pull down EGFP-LphD (or EGFP) and blotting for the binding of endogenous KAT7 we confirmed that KAT7 indeed interacts with endogenous EGFP-LphD, but not with EGFP alone (**Figure 4E**). Importantly, the binding of LphD to KAT7 is independent of its enzymatic activity, as catalytically inactive LphD-Y392F still binds KAT7 (**Figure 4E**, for IP control see **Figure S3C**). Reverse immunoprecipitation further verified that KAT7/LphD complex formation takes place in transfected cells (**Figure S3D**). Co-IP of EGFP-LphD and EGFP-RomA and further analyses of the presence of the different components of the HBO1 complex revealed that LphD interacts with all components of the HBO1 complex (BRPF1, KAT7, MEAF6 and ING5), whereas RomA binds only KAT7 (**Figure 4F**, for IP control see **Figure S3E**). Importantly, we could concomitantly show that RomA and LphD both bind the HBO1 complex, as well as the target histone H3.

To further analyse these interactions during infection, we used HEK293T cells that stably express the macrophage Fcg-RII receptor ^26^ which allows to engulf bacteria that have been opsonized with antibodies, leading to an efficient infection in combination with high transfection efficiency ^27^. We transfected these cells with either EGFP or EGFP-KAT7 and infected them using *L. pneumophila* wild type strain overexpressing V5-tagged forms of LphD or RomA. When pulling down the EGFP proteins we clearly detected an enrichment of both LphD and RomA in the EGFP-KAT7 sample (**Figure 4G**). *In vitro* binding assays with tagged forms of LphD, RomA and KAT7 produced in *E. coli* showed that RomA and LphD bind KAT7 also in a chromatin-free context **(Figure 4H** and **4I**), thus binding of LphD and RomA to KAT7 is specific and independent of chromatin targeting. To further explore the role of KAT7 during *L. pneumophila* infection and the synergistic activities of LphD and RomA, we quantified the levels of H3K14 methylation during infection with *L. pneumophila* wild type or the Δ*lphD* mutant strain by pre-treating the cells with a KAT7 specific inhibitor (WM-3835) (**Figure 2G**). This revealed that inhibition of KAT7 activity abolishes the decrease in H3K14 methylation observed in infection with *L. pneumophila* lacking LphD (Δ*lphD*). Thus, LphD counteracts KAT7 activity (**Figure 4J and Figure S3F**)

Taken together, these results further support our model that the two effectors, RomA and LphD, both target the KAT7 complex and chromatin to cooperatively regulate H3K14 acetylation and methylation for the benefit of intracellular growth of *L. pneumophila*.

## DISCUSSION

Analyses of the *L. pneumophila* genome identified a secreted nucleomodulin, named LphD, that posesses histone deacetylase activity. The study of the functional role of LphD in infection revealed the exceptional capacity of these bacteria to manipulate epigenetic marks of the host cell. Several bacterial effectors that target host chromatin have been identified in the last decade ^8^, however, the fact that different bacterial effectors may manipulate the host cell chromatin in synergy was unknown.

In eukaryotic cells, HDACs suppress gene expression by condensing chromatin packing and consequently decreasing chromatin accessibility for transcription factors and their binding to gene promoters ^28^. The regulation of their activity is a complex, multilayer process depending, among others, on the subcellular localization and protein complex formation. Although several bacterial pathogens have been shown to impact histone acetylation/deacetylation during infection by acting on expression of eukaryotic HDACs or their localization ^12^, until now only the *Neisseria gonorrhoeae* Gc-HDAC protein has been shown to induce an enrichment of H3K9ac, in particular at the promoters of pro-inflammatory genes ^29^.

Here we show that LphD encoded by *L. pneumophila* targets the host cell nucleus during infection, and deacetylates H3K14. The processes by which LphD targets the nucleus remains to be determined. Eukaryotic HDAC proteins are predominantly located in the nucleus, but some also shuttle between the nucleus and the cytoplasm, where they can also target cytoplasmic proteins. Class I HDACs whose subcellular localization is predominantly nuclear encode a nuclear localization sequence (NLS). Indeed, a putative NLS was predicted *in silico* at the N-terminus of LphD; however, its deletion did not affect nuclear translocation of LphD (**Figure S4A**). Thus, either another NLS is present in LphD that was not identified, or the nuclear import of LphD might be achieved by hijacking other HDAC-containing complexes such as NuRD. A key component of the NuRD complex, Retinoblastoma binding protein 4 (RBBP4), has been shown to directly interact with importin-a and to regulate the nuclear import ^30 31^. We identified several components of the NuRD complex as possible interaction partners of LphD, suggesting that LphD hijacks the NuRD complex to translocate to the host cell nucleus (**Table S1**), however, further experiments are necessary to confirm this hypothesis.

*In vitro* activities on short H3 peptides acetylated on different lysine residues showed a clear preference of LphD for H3K14 (**Figure 1F**), supported by an AlphaFold model of the complex between LphD and the histone H3 peptide tail, where H3K14 is placed in the active site pocket (**Figure 1E** and **inset**). Thus, H3K14 is the residue for which LphD exhibits the highest enzymatic efficiency, but the enzyme loses specificity if the reaction is allowed to proceed for a long time or when high concentrations of enzyme are present. This is seen in transfection when high amounts of protein are delivered into a host cell, as LphD transfection in HeLa cells leads to the deacetylation of several other residues such as H3K18 and H3K23 (**Figure S4B**). In contrast, during infection, when the enzyme is delivered in the host cell at physiological levels, only H3K14 is deacetylated (**Figures 2E and 2F**).

The deacetylation of H3K14 facilitates the methylation of the same residue by another bacterial effector, the SET-domain methyltransferase RomA. Importantly, we observed here that LphD and RomA not only target the same host protein, but that RomA activity is dependent on the presence of LphD as the level of H3K14 methylation during infection is significantly reduced in a Δ*lphD* background (**Figure 2G**). The cooperation between LphD and RomA was also observed in intracellular infection models and reflected in the transcriptome profiles (**Figure 3 and S3A**). The dependency of the two effectors on each other is even more striking in the phenotype of the double knockout. In replication assays, the Δ*lphD*Δ*romA* strain shows a phenotype closely resembling the wild type, compensating for the effect observed in the two single knockouts. Moreover, this compensation is reversible by complementation of either LphD or RomA, an effect not seen when using the catalytically inactive versions of the proteins for complementation analyses. This suggests that the enzymatic activities of LphD and RomA are only beneficial for the bacteria when both partners are present through a mechanism of regulation that is yet to be understood. Thus, RomA and LphD are “para-effectors”, from the Greek παρα(para) meaning *besides* but also *contrary to*, underlining the high interdependence of these two effectors. At least three *L. pneumophila* effectors have been described to target the same host protein, the small GTPase Rab1 and to sequentially modify it ^32^. However, uniquely we show here that the lack of the activity of one effector decreases the activity of the other one and both target the endogenous chromatin binding complex KAT7/HBO1 (**Figure 4**). Indeed, LphD and RomA immunoprecipitate host chromatin with an affinity for KAT7, as confirmed by *in vitro* binding assays (**Figure 4H** and **4I**). However, the fact that inhibition of KAT7 reverts the phenotype seen on H3K14me levels during Δ*lphD* infection might also imply that LphD directly modifies KAT7 acetylation. To explore the hypothesis that LphD also deacetylates KAT7 to modulate its activity, further experiments will be undertaken. Indeed, how KAT7 activity is regulated in a cell is still not known. Interestingly, KAT8, a close homologue of KAT7 regulates its activity through autoacetylation of a lysin residue that is also conserved in KAT7, also leaving the possibility that KAT7 regulation functions in the same manner ^33^ and suggesting that KAT7 might be a non-histone target of LphD.

LphD might have complex roles in the host cell and be implicated in regulating several histone and non-histone targets as the analysis of possible eukaryotic interaction partners of LphD led not only to the identification of KAT7/HBO1, but also suggested interaction with several other complexes related to chromatin remodeling. Some of these complexes, such as the NuRD-and the Sin3-complex, are known to comprise eukaryotic histone deacetylases and might thus be additional interaction partners for LphD ^34 35^. Further studies will elucidate whether additional interaction partners of LphD and RomA exist, that could be important during infection of different host cells, as *L. pneumophila* is known to infect many protozoan hosts and different mammalian cells. In conclusion, this study provides exciting insight into how *L. pneumophila* modifies host chromatin by using two distinct chromatin remodelers that function in synergy. Both LphD and RomA are deployed by the bacteria to strategically influence the response of the host cell to infection and promote bacterial replication in this otherwise hostile environment.

## MATERIALS and METHODS

### Bacterial strains, growth conditions and cell culture

*Legionella pneumophila* strain Paris and mutants were cultured in N-(2-acetamido)-2-aminoethanesulfonic acid (ACES)-buffered yeast extract broth (BYE) or on ACES-buffered charcoal-yeast (BCYE) extract agar ^36^. For *Escherichia coli* Luria-Bertani broth (LB) was used. When needed antibiotics were added: for *L. pneumophila (E. coli):* kanamycin 12.5 μg/ml (50 μg/ml), gentamycin 12.5 μg/ml, apramycin 15 μg/ml, chloramphenicol 10 μg/ml (10 μg/ml), and ampicillin (only for *E. coli*) 100 μg/ml.

THP-1 cells were maintained in RPMI-1640 (Gibco), HeLa and HEK293T in DMEM GlutaMAX (Gibco), both containing 10% FBS (Eurobio Scientific) in a humid environment with 5% CO_2_ at 37°C. *Acanthamoeba castellanii* [ATCC50739] was cultured at 20°C in PYG 712 medium [2% proteose peptone, 0.1% yeast extract, 0.1 M glucose, 4 mM MgSO_4_, 0.4 M CaCl_2_, 0.1% sodium citrate dihydrate, 0.05 mM Fe(NH_4_)_2_(SO_4_)_2_·6H_2_O, 2.5 mM NaH_2_PO_3_, 2.5 mM K_2_HPO_3_]. Cell transfections were performed by using FuGENE (Promega) following the recommendations of the manufacturer.

### Constructions of mutants and complementation plasmids

The knockout of *lphD* in the wild type background to generate a single mutant followed by the knockout of *romA* to generate the double mutant, was performed as previously described ^37^. To construct the Δ*lphD* mutant strain the chromosomal gene *lphD* of the wild type strain was replaced by introducing a gentamycin resistance cassette. The mutant allele was constructed using a 3-steps PCR. Briefly, three overlapping fragments (*lphD* upstream region-primers 195H and 196H, antibiotic cassette-primers 52H and 52B, *lphD* downstream region-primers 195B and 196B; **Table S2**) were amplified independently and purified on agarose gels. The three resulting PCR products were mixed at the same molar concentration (15nM) and a second PCR with flanking primer pairs (primers 195H and 195B; **Table S2**) was performed. The resulting PCR product, the gentamycin resistance cassette flanked by 500 bp regions homologous to *lphD* was introduced into strain *L. pneumophila* Paris by natural competence for chromosomal recombination. Strains that had undergone allelic exchange were selected by plating on BCYE containing gentamycin and the mutant was verified by PCR and sequencing. The resulting mutant was then used to generate the Δ*lphD ΔromA* double knockout strain, using a new set of primers (*romA* upstream region-primers 11H and 66H, kanamycin antibiotic resistance cassette-primers 60H and 60B, *romA* downstream region-primers 11B and 66B; **Table S2**). The generated mutant strains were whole genome sequenced to confirm the correct deletion of the gene and the absence of other mutations in the genome.

For the complementation construction, the full-length *lphD* with its own promotor was cloned into pBC-KS (Stratagene). Bacteria expressing EGFP were obtained by introducing EGFP under the control of the *flaA* promotor of *L. pneumophila* into pBC-KS backbone. To generate the catalytic inactive Y392F mutant of LphD, a single base pair mutation was performed using mismatched primers (217H and 217B, **Table S2**).

### β-lactamase translocation assays

β-lactamase assays were performed in THP1 infected cells as previously described ^38^. Around 10^5^ THP-1 cells are seeded in a 96-well plate and differentiated for 72 hours using 10 nM phorbol 12-myristate 13-acetate (PMA). One day before infection *L. pneumophila* wild type or <u>Δ*dotA* mutant strain</u> carrying plasmids for the expression of either β-lactamase alone or a β-lactamase-LphD or -RomA fusion are cultured in BYE broth containing chloramphenicol and IPTG (1 mM) to induce protein production. After differentiation the cells are washed, fresh RPMI medium with 1mM IPTG is added. Cells are infected with the β-lactamase fusion protein expressing bacteria at an MOI of 50. Spin the plates for 5 minutes at 300 g and then incubate the plate for 2 hours at 37°C in a humidified 5% CO_2_ atmosphere. After this incubation, LiveBLAzer CCF4-AM solution (Thermo Fisher Scientific) is added to all the wells and the plate is incubated in the dark at room temperature for 2 hours. The cells are washed, and cell dissociation solution (SIGMA) is added. After another incubation of 30 minutes at 37°C with *5%* CO_2_ the samples are analysed by flow cytometry (MACS Quant, Miltenyi Biotec) and FlowJo™ v10.8 Software (BD Life Sciences). Non-infected unstained cells are used as negative control. Stained uninfected cells were used to gate the green positive cells (uncleaved CCF4). For gating strategy see **Figure S2A**.

### Infection Assays

*A. castellanii* were washed once with infection buffer (PYG 712 medium without proteose peptone, glucose and yeast extract) and seeded at a concentration of 4×10^6^ cells per T25 flask. *L. pneumophila* wild type and mutant strains were grown on BCYE agar to stationary phase, diluted in infection buffer and mixed with *A. castellanii* at an MOI of 0.1 or 1 (for complementation assays). Infected cells were maintained at 20°C and intracellular multiplication was monitored plating a sample at different time points (2 h, 24 h, 48 h, 72 h, 96 h, 120 h, 144 h, and 168h) on BCYE plates and the number of intracellular bacteria was counted. In THP-1 cell infection assays 10^6^ cells were split in conical tubes (Falcon, BD lab ware). Stationary phase *L. pneumophila* were resuspended in serum free medium and added to cells at an MOI of 10. After 1 hour of incubation, Gentamycin (100 μg/ml) was added. After another hour of incubation, the infected cells were washed with PBS, before incubation with 2 ml of RPMI. At 2 h, 24 h and 48 h, 500 μl of cell suspension were mixed with equal amounts of PBS-0.2% TritonX-100 for lysis. The infection efficiency was monitored by determining the number of colony-forming units (cfu) of the different *L. pneumophila* strains after plating on BCYE agar.

### Immunofluorescence analyses

For immunofluorescence analyses, cells are fixed with PBS-4% para-formaldehyde for 15 minutes at room temperature, followed by quenching (PBS-50 mM NH4Cl) for 10 minutes. Cells are permeabilized with PBS-0.1% Triton X-100 and blocked for 30 minutes with PBS-5% BSA. The cells are incubated with the respective primary antibodies overnight at 4°C in PBS-5% BSA. They are washed three times using PBS and then stained with DAPI, phalloidin and secondary antibodies for 30 minutes at room temperature, followed by mounting to glass slides using Mowiol (SIGMA). Antibodies used in this study are listed in **Table S3**. Immunosignals were analyzed with a Leica SP8 Microscope at 63× magnification. Images were processed using Fiji software ^39^.

### LphD purification

N-terminal HIS_6_-tagged LphD was expressed in *E. coli* BL21 C41 following an auto-induction protocol ^40^. After 4 hours at 37°C cells were grown for 20 hours at 20°C in 2YT complemented autoinduction medium containing 50 μg/ml kanamycin. Cells were harvested and flash frozen in liquid nitrogen. Cell pellets were resuspended in 50ml lysis buffer (50 mM HEPES pH8, 500 mM KCl, 5% glycerol, 1 mM MgCl2, benzonase, lysozyme, 1 mM DTT and supplemented with EDTA-free protease inhibitor cocktails (ROCHE)) at 4°C and disrupted by sonication (6 x 60 seconds). The lysate was centrifuged for 60 min at 10,000 g at 4°C. The cleared lysate was loaded onto a Ni-NTA affinity chromatography column (HisTrap FF crude, GE Healthcare) equilibrated in Buffer A (50 mM Hepes pH8, 500 mM KCl, 5% glycerol, 10 mM imidazole, 1 mM DTT). HIS_6_-tagged proteins were eluted with a linear gradient of buffer B (50 mM Hepes pH8, 500 mM KCl, 5% glycerol, 1 M imidazole, 1 mM DTT). The eluted fractions containing the protein of interest were pooled and dialysed at 4°C overnight in SEC buffer (20 mM HEPES pH8, 300 mM KCl, 5% glycerol, 2 mM TCEP). The HIS_6_-tag was not removed as this led to precipitation of the protein. After dialysis, the protein was concentrated and loaded onto a Superdex 75 16/60 size exclusion (SEC) column (GE Healthcare). The peak corresponding to the protein was concentrated to about 12 mg/ml and flash frozen in liquid nitrogen and stored at −80°C.

### UFLC-mediated LphD deacetylase activity assay

In order to quantify LphD deacetylase activity, we synthetized six 5-fluorescein amidite (5-FAM)-conjugated acetylated peptide substrates based on the human H3.1 sequence and centered on various lysine residues of interest:

ARTK_ac_QTARRSK-(5-FAM), referred to as H3K4ac peptide; (5-FAM)-QTARK_ac_STGG-NH_2_, referred to as H3K9ac peptide; (5-FAM)-STGGK_ac_APRR-NH_2_, referred to as H3K14ac peptide; (5-FAM)-RAPRK_ac_QLAT-NH_2_, referred to as H3K18ac peptide; (5-FAM)-QLATK_ac_AARR-NH_2_, referred to as H3K23ac peptide; (5-FAM)-TRAARKacSAPAT-NH2, referred to as H3K27ac peptide. Non-acetylated versions of these peptides were also synthetized as detection standards. Samples containing H3 peptides, and their acetylated forms were separated by RP-UFLC (Shimadzu) using Shim-pack XR-ODS column 2.0 x 100 mm 12 nm pores at 40°C. The mobile phase used for the separation consisted of the mix of 2 solvents: A was water with 0.12% trifluoroacetic acid (TFA) and B was acetonitrile with 0.12% TFA. Separation was performed by an isocratic flow depending on the peptide: 83 % A/17 % B, rate of 1 ml/min, 6 min run for H3K4ac peptide; 80 % A/20 % B, rate of 1 ml/min, 6 min run for H3K9ac, H3K14ac, H3K18ac, H3K27ac peptides; 79 % A/21 % B, rate of 1 ml/min, 8 min run for H3K23ac peptide. H3 acetylated peptides (substrates) and their non-acetylated forms (products) were monitored by fluorescence emission (λ = 530 nm) after excitation at λ = 485 nm and quantified by integration of the peak absorbance area, employing a calibration curve established with various known concentrations of peptides. The kinetic parameters of LphD on H3-derived peptides were determined by UFL in a 96-wells ELISA. Briefly, LphD (7.7 nM) was mixed with different concentrations of acetylated H3 peptides (ranging from 12.5 to 200 *μ*M final) for 15 minutes at 30°C and the reaction was stopped by adding 50 *μ*L of HClO_4_(15% v/v in water). Finally, 10 *μ*l of the reaction mix were automatically injected into the RP-UFLC column and initial velocities (V_i_, *μ*M.min^-1^) were determined as described above. V_i_ were then plotted against substrate peptide concentrations and curves were non-linearly fitted using Michaelis–Menten equation 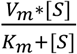 (OriginPro 8.0). K_m_ (enzyme Michaelis’s constant), V_m_ (enzyme maximal initial velocity) and k_cat_ (enzyme turnover) values were extrapolated from these fits. A catalytic dead version of the enzyme was used as a negative deacetylation control^41^.

### Highly acetylated histone extraction

HEK293T cells were cultivated in RPMI 1640 medium with 10 % heat-inactivated fetal bovine serum (FBS) and 1 mM L-glutamine at 37°C under 5 % CO_2_. For endogenous histone extraction, cells were seeded at 30000 cells/cm^2^ in a 100 cm^2^ Petri dish (VWR). The next day, cells were treated with 20 mM sodium butyrate and 6 μM Trichostatin A (TSA). Cells were then put back in the incubator at 37°C and 5 % CO_2_ for 30 min before being harvested. Cells were lysed with cell lysis buffer (PBS 1x, 1% Triton X-100, 20 mM sodium butyrate, 6 μM TSA, protease inhibitors) for 30 min at 4°C, sonicated (2 sec, 10 % power) and centrifuged (15 min, 15500 g, 4°C). 500 *μ*L of 0.2 N HCl was then put on remaining pellets. The mixture was sonicated 3 times (3 sec, 10 % power) and incubated overnight at 4°C. The next day, samples were centrifuged (15 min, 15500 g, 4°C) and the supernatant (containing extracted histones) was buffer exchanged three times into Tris 50 mM, 50 mM NaCl, pH 8 using MiniTrap G-25 desalting columns (GE Healthcare) and stored with protease inhibitor at −20°C until use.

### *In vitro* histone deacetylase activity assay

Assays were performed in a total volume of 20 *μ*L LphD buffer containing 250 ng highly acetylated, purified histones and 10 ng LphD (wild type or catalytic inactive mutant LphD Y392F) at 30°C. At different time points (0, 1, 2, 3, 4 and 5 minutes), the reaction was stopped with the addition of 10 *μ*L Laemmli sample buffer containing 400 mM β-mercaptoethanol.

Samples were analyzed on 18% SDS PAGE, followed by a transfer onto a nitrocellulose membrane (0.2 *μ*m) at 200 mA for 65 min. Ponceau staining was carried out to ensure equal protein loading. Membranes were blocked with non-fat milk (5%) in PBS-1% Tween (PBST) for 1 h and incubated with α-H3K14Ac antibody in 1% non-fat milk PBST overnight at 4°C. After washing 3 times, the membranes were incubated for 1 h at room temperature with peroxidase-coupled secondary antibody in 1% non-fat milk PBST. The proteins were then visualized by chemiluminescence detection using ECL reagent on LAS 4000 (Fujifilm) instrument. Images were processed and quantified using MultiGauge V3.0 and Fiji software.

### Fluor de Lys^®^ *in vitro* enzymatic assays

Purified HIS_6_-LphD was used to perform *in vitro* enzymatic assays with Fluor de Lys^®^deacetylase assay (Enzo Life Sciences). Briefly, different amount of purified LphD and catalytic inactive mutant LphD Y392F were incubated with the substrate in a reaction buffer (20 ml; 50mM TRIS/Cl, pH 8.0, 137mM NaCl, 2.7mM KCl, 1mM MgCl2) for 30 minutes at 37°C. The signal was then read using a plate reader (TECAN). Trichostatin-A (TSA) was added at 5 *μ*M.

### Histone modification analysis of infected cells

For the analysis of histone modifications during infection, THP-1 cells were infected at an MOI of 50 with *L. pneumophila* wild type or mutant strains, both containing a plasmid for the expression of EGFP under the control of the *flaA* promoter. After 30 minutes, Gentamicin is added (100 *μ*g/ml) to kill extracellular bacteria. Cells are then sorted by FACS (S3e, BIORAD) as previously described ^42^. Histones of infected cells were isolated as previously described with some modifications ^43^. Briefly, THP-1 cells (3×10^6^) were incubated at 4°C with hypotonic lysis buffer (10 mM Tris–HCl pH 8.0, 1 mM KCl, 1.5 mM MgCl_2_, with protease inhibitors (ROCHE)) for 2 hours while rotating. Subsequently, nuclei were pelleted and resuspended in 0.4 M sulfuric acid and incubated overnight at 4°C. The supernatant was precipitated with 33% trichloroacetic acid (TCA) on ice. Pelleted histones were washed twice with ice-cold acetone and were then resuspended in DNase/RNase free water. Sample quality of acid extraction was visualized on a Coomassie-stained 4-15% SDS-PAGE. Histone modification signal (H3K14ac, H3K14me, H3K18ac, H3K23ac) is assessed by immunoblot and normalized to signal of histone H1. Sample signals are then compared to non-infected controls.

### Co-immunoprecipitation experiments

For the GFP-pulldown HEK293T cells were seeded in 10 cm dishes and transfected 24 or 48 hours with 3 *μ*g of the different EGFP construct expression plasmids. Transfected cells were washed three times with PBS before lysis in RIPA buffer (20 mM HEPES-HCl pH 7.4, 150 mM NaCl, 5 mM EDTA, 1% Triton X-100, 0.1% SDS, with protease inhibitors (ROCHE)). For the verification of protein interaction during infection we modified the previously established protocol ^27^. Briefly, we transiently transfected HEK293-FcgRII cells with either EGFP or a EGFP-KAT7 fusion product. After 48 hours of transfection, the cells were washed and fresh DMEM with IPTG (1 mM) was added. *L. pneumophila* over-expressing either V5-LphD or V5-RomA were grown until post exponential phase in presence of 1mM IPTG to allow fusion protein expression. After reaching an OD ~4, the bacteria were pre-opsonized by incubating them with an anti-FlaA antibody for 30 minutes at 37°C. Then the cells are infected with MOI of 50. After one hour, the cells are washed and fresh DMEM (with 1 mM IPTG) is added. After 7 hours of infection, the cells are collected and lysed in RIPA buffer. To facilitate the lysis, cells were sonicated using a Bioruptor^®^ Pico sonication device (Diagenode) for 15 cycles of 30 seconds ON/OFF. Lysates were precleared and the pulldown was performed using GFP-trap magnetic agarose beads (Chromotek), following the manufacturer’s instructions at 4°C overnight. Proteins were eluted in 30 *μ*l Laemmli buffer and then analyzed by western blot or the beads directly processed for MS/MS analyses.

### Mass spectrometry analysis of GFP co-IP

MS grade Acetonitrile (ACN), MS grade H_2_O and MS grade formic acid (FA) were acquired from Thermo Chemical. Proteins on magnetic beads were digested overnight at 37°C with 1_μ_l (0.2 _μ_g/_μ_L) of trypsin (Promega) in a 25-mM NH_4_HCO_3_ buffer per sample. The resulting peptides were desalted using ZipTip μ-C18 Pipette Tips (Pierce Biotechnology). Samples were analyzed using an Orbitrap Fusion equipped with an easy spray ion source and coupled to a nano-LC Proxeon 1200 (Thermo Fisher Scientific). Peptides were loaded with an online preconcentration method and separated by chromatography using a Pepmap-RSLC C18 column (0.75 × 750 mm, 2 μm, 100 Å) from Thermo Fisher Scientific, equilibrated at 50°C and operating at a flow rate of 300 nl/min. Peptides were eluted by a gradient of solvent A (H_2_O, 0.1 % FA) and solvent B (ACN/H_2_O 80/20, 0.1% FA), the column was first equilibrated 5 min with 95 % of A, then B was raised to 28 % in 105 min and to 40% in 15 min. Finally, the column was washed with 95% B during 20 min and re-equilibrated at 95% A during 10 min. Peptide masses were analyzed in the Orbitrap cell in full ion scan mode, at a resolution of 120,000, a mass range of *m/z* 350-1550 and an AGC target of 4.10^5^. MS/MS were performed in the top speed 3s mode. Peptides were selected for fragmentation by Higher-energy C-trap Dissociation (HCD) with a Normalized Collisional Energy of 27% and a dynamic exclusion of 60 seconds. Fragment masses were measured in an Ion trap in the rapid mode, with and an AGC target of 1×10^4^. Monocharged peptides and unassigned charge states were excluded from the MS/MS acquisition. The maximum ion accumulation times were set to 100 ms for MS and 35 ms for MS/MS acquisitions, respectively. Label-free quantification was done on Progenesis QI for Proteomics (Waters) in Hi-3 mode for protein abundance calculation. MGF peak files from Progenesis were processed by Proteome Discoverer 2.4 with the Sequest search engine. A custom database was created using the Swissprot/TrEMBL protein database release 2019_08 with the *Homo sapiens* taxonomy and including LphD, both from the *Legionella pneumophila* taxonomy. A maximum of 2 missed cleavages was authorized. Precursor and fragment mass tolerances were set to respectively 7 ppm and 0.5 Da. Spectra were filtered using a 1% FDR using the percolator node.

### *In vitro* protein binding assays

To assess protein-protein interactions *in vitro*, purified GST-RomA was incubated with equal amounts of HIS_6_-LphD or HIS_6_-LphD Y392F in protein binding buffer (25 mM Tris pH 8.0, 140 mM NaCl, 3 mM KCl, 0.1% NP40 with protease inhibitors (ROCHE)) overnight at 4°C. As binding control, the LphD proteins were also incubated with GST-MLY1 (myosin light chain 1/3) under the same conditions. Then GST-tagged proteins were pulled down using Glutathione magnetic agarose (Jena Bioscience). Beads were washed with protein binding buffer and proteins are then eluted using Laemmli buffer. Then samples were analysed by immunoblot. For the interaction of KAT7 with LphD and RomA the same approach was used. FLAG-KAT7 was incubated with either HIS6-LphD or GST-RomA (HIS6-MBP or GST-MLY1 as binding controls) and the pulldown was performed using DYKDDDDK Fab-Trap beads (Chromotek) followed by western blot analysis.

### Western blotting

Sample proteins were prepared in Laemmli sample buffer containing 400 mM β-mercaptoethanol and loaded on SDS PAGE gels, followed by a transfer onto a nitrocellulose membrane (0.2 *μ*m; Trans-Blot Turbo system, Biorad). Ponceau S (SIGMA) staining was carried out to ensure equal protein loading. Membranes were blocked with 5% non-fat milk in TBS-Tween 0.5% for 1 hour and incubated with the respective primary antibody overnight at 4°C. Antibodies used are listed in **Table S3**. Membranes were washed and probed with horseradish peroxidase-coupled antibody against either mouse IgG or rabbit IgG (1:2500 in 5% non-fat milk TBS-Tween) for 1 hour. The proteins were then visualized by chemiluminescence detection using HRP Substrate spray reagent (Advansta) on the G:BOX instrument (Syngene). Images were processed and quantified using Fiji software ^39^.

### RNA-sequencing

For the RNA-seq THP-1 monocytes were infected with either *L. pneumophila* wild type, Δ*lphD*, Δ*romA* or Δ*lphD*-Δ*romA* strains, all of which were expressing EGFP under control of the *flaA* promoter. After 7 hours post-infection, the infected cells were enriched by FACS and stored at −80°C. RNA-seq and data analysis was performed by Active Motif. In short, total RNA was isolated from samples using the Qiagen RNeasy Mini Kit (Qiagen). For each sample, 1 *μ*g of total RNA was then used in the TruSeq Stranded mRNA Library kit (Illumina). Libraries were sequenced on Illumina NextSeq 500 as paired-end 42-nt reads. Sequence reads were analyzed with the STAR alignment – DESeq2 software pipeline.

### ChIP Experiments

THP-1 monocytes were infected with either *L. pneumophila* wild type or Δ*lphD* strain, which both were expressing EGFP under control of the *flaA* promoter. After 7 h post-infection, the infected cells were enriched by FACS and stored at −80°C until chromatin isolation. Chromatin immunoprecipitation (ChIP) experiments were undertaken as previously described ^20^. DNA enrichment was followed by quantitative PCR (qPCR) (QuantStudio Thermo Fisher Scientific) with the primers listed in **Table S2** and normalized using the Percent Input Method (ChIP signals divided by input sample signals).

### Production of anti-LphD antibodies in rabbit

Rabbit polyclonal antibodies to LphD were generated for this study (Thermo Fisher). Briefly, the purified His6-LphD was injected in a rabbit for a protocol of 90 days. The produced antibody was qualitatively evaluated by affinity purified ELISA: the purified antibody is tittered by indirect ELISA against the protein bound to the solid-phase to measure the reactivity of the antibodies after elution.

## Supporting information

Supplemntal data for Schator et al

## DATA AVAILABILITY STATEMENT

The sequence reads as well as the coverage files of the RNAseq libraries of THP-1 cells have been deposited in the NCBI Gene Expression Omnibus (GEO) database ^44^. Accession number GSE207487. Source data are provided with this paper.

## AUTHOR CONTRIBUTIONS

D.S., M.R. and C.B. designed research; D.S., M.R., S.M., J.B., C.D.S. C.R. and M.B.A. performed experiments; F.R.L. C.R. D.S. MR. S.M. and A.W. analysed data; and D.S., M.R. and C.B. wrote the paper.

## ACKNOWLEDGMENTS

Work in the C.B. laboratory is financed by the Institut Pasteur, the Fondation pour la Recherche Médicale (FRM) grant N° EQU201903007847 and the Agence Nationale de la Recherche grant n°ANR-10-LABX-62-IBEID to C.B. and grant n° ANR-18-CE15-0005-01 to M.R. D.S. was funded by a Sorbonne University doctoral contract. CDS is funded by an “Université de Paris Cité” doctoral contract. We thank the members of the ProteoSeine@IJM facility (Institut Jacques Monod, CNRS UMR 7592 Paris University and the region Île-de-France) for mass spectrometry analyses. We thank Giulia Brenna for technical help and the group of Craig Roy for providing the HEK293-FcγRII cells. We gratefully acknowledge the UtechS Photonic BioImaging, C2RT, Institut Pasteur, supported by the French National Research Agency (France BioImaging; ANR-10–INSB–04; Investments for the Future) and the technical platform “BioProfiler-UFLC” (BFA Unit, Université Paris Cité) for provision of UFLC facilities.

