## Supplementary material for "*Legionella* para-effectors target chromatin and promote bacterial replication": Supplemntal data for Schator et al

#### Materials and Methods

**Table S1:** LphD interacting proteins identified through GFP-trap analysis that are known to be involved in epigenetic regulation

**Table S2:** Primers used in this study

**Table S3:** Antibodies and dyes used in this study

**Figure S1:** LphD Alignment H3/H4 peptide accommodation and activity on octamers.

**Figure S2:** Anti-LphD validation, representative immunoblots for histone deacetylation and complementation assays.

**Figure S3:** IP controls and Immunoblots showing the interaction of LphD with KAT7

**Figure S4**: Subcellular localisation of LphD deleted of its putative NLS and H3K18ac/H3K23ac levels upon LphD transfection.

### SUPPLEMENTARY TABLES

**Table S4:** LphD interacting proteins identified through GFP-trap analysis that are known to be involved in epigenetic regulation

| Protein | Epigenetic complex | FDR | Fold change |
|---|---|---|---|
| KAT7 | HBO1 | 0.06 | 75.70 |
| HDAC2 | NuRD/Sin3 | 0.08 | 8.44 |
| MTA2 | NuRD | 0.03 | 5.41 |
| SAP18 | Sin3 | 0.005 | 9.46 |
| PRKDC | DNA-PK | 0.005 | 4.71 |
| XRCC5 | DNA-PK | 0.02 | 14.47 |
| SSRP1 | FACT | 0.02 | 3.08 |
| SUPT16H | FACT | 0.001 | 3.39 |
| EED | PRC2/EED-EZH2 | 0.03 | 32.19 |
| SUZ12 | PRC2/EED-EZH2 | 0.002 | 15.45 |
| EZH2 | PRC2/EED-EZH2 | 0.09 | 9.17 |

**Table S5:** Primers used in this study

| Primer | Sequence (5'-3') | Purpose | Reference |
|---|---|---|---|
| 52H | GATGAAGGCACGAACCCAGTTGACA | Deletion of *lphD* gene | This study |
| 52B | CGGCTTGAACGAATTGTTAGGTGGC | Deletion of *lphD* gene | This study |
| 195H | GACCCTCGACTTAATTGGATAACGG | Deletion of *lphD* gene | This study |
| 195B | GGAGGGTAAACGGAAAACAAACTG | Deletion of *lphD* gene | This study |
| 196H | GCCACCTAACAATTCGTTCAAGCCGGGTGATC TAAGCTACTTCTATGAATCCATATTC | Deletion of *lphD* gene | This study |
| 196B | TGTCAACTGGGTTCGTGCCTTCATCGTTTCCTG AGTTGTAATAATTGGTAAATTCC | Deletion of *lphD* gene | This study |
| 11H | TCCAATATACAAGCATTCATGTGCTATCTG | Deletion of *romA* gene | [1] |
| 11B | GAAGTTTCTCGAATTCTTTGGACAAGC | Deletion of *romA* gene | [1] |
| 60H | CATCGATGAATTGTGTCTCAAAA | Deletion of *romA* gene | [1] |
| 60B | GTCCCGTCAAGTCAGCGTA | Deletion of *romA* gene | [1] |
| 66H | TTTTGAGACACAATTCATCGATG GCTCTATTT TGCATGTGATTTTCATT | Deletion of *romA* gene | [1] |
| 66B | TACGCTGACTTGACGGGAC GCAAGTTTTTTG ATTTGATATTTCCTG | Deletion of *romA* gene | [1] |
| 217H | ATCTATTGGGGCTTGGAAGGTGGATTTGACAG GACCATGTATGAA | Base pair substitution (LphD Y392F) | This study |
| 217B | TTCATACATGGTCCTGTCAAATCCACCTTCCAA GCCCCAATAGAT | Base pair substitution (LphD Y392F) | This study |
| 222H | GGATCCGAAAGTAGCGCCTCTTGTCTTGATA | Complementation of *lphD* | This study |
| 222B | GGTACCTTAACAGGACATATTATGCGAATTGG | Complementation of *lphD* | This study |
| CP_07H | CTCTCAATCTCCAGCCACAAAT | CXCL1 qPCR ChIP | [1] |
| CP_07B | CCTGAGAACCACCACAGAGAAG | CXCL1 qPCR ChIP | [1] |
| CP_26H | GCTCCCCAACCTAGTGTCAT | MyD88 qPCR ChIP | [1] |
| CP_26B | GGAGTGGGAAACGGACAG | MyD88 qPCR ChIP | [1] |
| CP_37H | AGTTGTGGTCTGTGGCACTC | IFNB1 qPCR ChIP | [1] |
| CP_37B | AGTTTGGGCTTTCTCACAGC | IFNB1 qPCR ChIP | [1] |
| CP_30H | GATGTGGAGCTGGGATGTC | BCL2L1 qPCR ChIP | [1] |
| CP_30B | CATGGCAGCAGTAAAGCAAG | BCL2L1 qPCR ChIP | [1] |
| CP_22H | TGAGAAAGGAGGTGGGTAGG | IL6 qPCR ChIP | [1] |
| CP_22B | CCCAGCAAAGACCTCCTAAT | IL6 qPCR ChIP | [1] |
| AJUBA_fwd | CCAGGAATCCCACAGCATCC | AJUBA qPCR ChIP | [1] |
| AJUBA_rev | CGGGGAGAGTGGGGCA | AJUBA qPCR ChIP | [1] |
| TXNIP_fwd | ATTGGGCCGCTTACCTGTTG | TXNIP qPCR ChIP | [1] |
| TXNIP_rev | GTTAGGATCCTGGCTTGCGG | TXNIP qPCR ChIP | [1] |

**Table S6:** Antibodies and dyes used in this study

| Target | Manufacturer | Product code | Dilution |
|---|---|---|---|
| Western blot: | | | |
| b-Actin | Sigma | A5316 | 10.000 |
| BRPF1 | Active Motif | 61541 | 1000 |
| EGFP | Thermo Fisher | A11122 | 2000 |
| FLAG | Sigma | F3165 | 2000 |
| GST | Milipore | AB3282 | 1000 |
| H1 | Active Motif | 61201 | 2000 |
| H3 | Active Motif | 39163 | 2000 |
| H3K14ac | Milipore | 07-353 | 2000 |
| H3K14me2 | Euromedex | H3-2B10 | 3000 |
| H3K18ac | Active Motif | 39755 | 1000 |
| H3K23ac | Active Motif | 39131 | 1000 |
| HA | Sigma | H6908 | 5000 |
| HDAC1 | Santa Cruz Biotechnology | sc-81598 | 500 |
| HIS | Sigma | H1029 | 2000 |
| ING5 | Active Motif | 91329 | 500 |
| KAT7 | Santa Cruz Biotechnology | sc-39846 | 500 |
| MEAF6 | Thermo Fisher | PA5-40704 | 1000 |
| V5 | Thermo Fisher | 46-0705 | 3000 |
| anti-Mouse HRP-conjugated | Cell signaling | 7076S | 2500 |
| anti-Rabbit HRP-conjugated | Cell signaling | 7074S | 2500 |
| Immunofluorescence: | | | |
| LphD | Thermo Fisher | Custom (see **Methods**) | 500 |
| Alexa488 goat anti-Rabbit | Thermo Fisher | A32731 | 1000 |
| Alexa546 goat anti-Rabbit | Thermo Fisher | A11010 | 1000 |
| DAPI | Thermo Fisher | D21490 | 600 |
| Alexa633 phalloidin | Thermo Fisher | A22284 | 500 |

#### SUPPLEMENTARY FIGURES LEGENDS

**Supplementary Figure S1: LphD Alignment H3/H4 peptide accommodation and activity on octamers.** (**S1A**) Multiple sequence alignment of LphD (aa 160-424) and several eukaryotic HDAC domains (Accession numbers: *Homo sapiens*: HDAC1 #Q6IT96, HDAC2 #Q92769, HDAC3 #O15379, HDAC4 #P56524, HDAC5 #Q9UQL6, HDAC6.1 #Q9UBN7, HDAC6.2 #Q9UBN7, HDAC7 #Q8WUI4, HDAC8 #Q9BY41, HDAC9 #Q9UKV0, HDAC10 #Q969S8, HDAC11 #Q96DB2; *Saccharomyces cerevisiae*: Rpd3 # P32561, HDA1 #P53973). Analysis also includes two yeast HDACs (Rpd3 and HDA1), which are the basis of general HDAC classification. Alignment was performed using SeaView and visualization with ESPript. Red boxes with white characters mean strict identity between all samples, bold characters mean high in-group similarity and yellow boxes mean high across-group similarity. Binding pocket residues (charge relay system) and the catalytic center are marked with black frames. (**S1B**) Histone tail residues identified as possible targets of LphD by MS/MS. Purified histones were incubated with or without purified LphD for 1 hour at 37°C, followed by MS/MS analysis. (**S1C**) Cartoon representation of the LphD-histone tail (H3) model generated by AlphaFold[2]. The per-residue confidence score (pLDDT) produced by AlphaFold is shown in the insert. Superposition of the 5 models of the LphD-H3 peptide complex. All 5 models place the same lysine 14 into the active site pocket. (**S1D**) Cartoon representation of the LphD-histone tail models (H4) generated by AlphaFold. Individual models of LphD-H4 complexes showing different binding modes for the H4 peptide. Only the first two place a lysine residue into the active site. (**S1E**) Validation of H3K14ac antibody specificity by dot blot experiments, using different amounts of several H3 peptides (H3K14, H3K14me, H3K14ac, H3K18, H3K18me, H3K18ac, H3K27, H3K27me, H3K27ac).

**Supplementary Figure S2: Anti-LphD validation, representative immunoblots for histone deacetylation and complementation assays.** (**S2A**) FACS Strategy of ß-lactamase secretion assay. Representative image of using FSC vs SSC plot to gate on live cells. THP1 cells were infected, as indicted, and after CCF4 loading samples were analyzed by flow cytometry to determine green (Y axis) and blue (X axis) cell populations, where green represents no translocation and blue represents translocation of the ß-lactamase into the cytosol. Double negative (DNeg; uninfected unstained) cells were used as negative control. Stained uninfected cells were used to gate the green positive cells (uncleaved CCF4). Representative gates of the analyzed conditions: ß-lac (uncleaved CCF4); ß-lac-RomA (positive control: cleaved CCF4 inducing a cell shift in blue channel); ß-LphD (as for the positive control: cleaved CCF4 and blue shift); ß-LphD in a *ΔdotA* strain (uncleaved CCF4). (**S2B**) Immunofluorescence analysis of subcellular localization of EGFP. HeLa cells were transfected with EGFP. DAPI (cyan), EGFP (green), and phalloidin (gray). Scale bars 10 $\mu$m. (**S2C**) Validation of custom anti-LphD antibody produced in rabbit. **Left**: Titer determination of polyclonal antibodies by ELISA comparing binding activity to purified $HIS_6$-LphD, compared to $HIS_6$-MBP. **Right**: Specificity testing by western blot using anti-LphD antibody. Line 1: purified $HIS_6$-LphD, Line 2: purified $HIS_6$-MBP, Line 3: *L. pneumophila* extract overexpressing V5-LphD. (**S2D, E F, G**) Representative image of western blots for H3K14ac (**S2D**), H3K18ac (**S2E**), H3K23ac (**S2F**), and H3K14me (**S2G**). THP-1 cells were infected with *L.pneumophila* wild type or *ΔlphD* strain expressing EGFP. Cells were sorted at different times post-infection by FACS, histones were isolated and analyzed by western blot. Histone H1 was used as loading control.

**Supplemental Figure S3: IP controls and Immunoblots showing the interaction of LphD with KAT7.** (**S3A**) Heat map of RNAseq results. Genes significantly up- ($\log_2$ fold change ≥ 2, p.adjust < 0.1) or down-regulated ($\log_2$ fold change ≥ -2, p.adjust < 0.1) (~4000 genes) during wild type *L. pneumophila* infection are analyzed and the impact of the three knockout strains (*ΔlphD*, *ΔromA* and *ΔlphDΔromA*) on these genes during infection is shown in terms of $\log_2$ fold change. THP-1 cells were infected with *L. pneumophila* wild type, *ΔlphD*, *ΔromA* or *ΔlphDΔromA* strains expressing EGFP. Cells were sorted 7 hours post-infection by FACS and RNA processed for the RNAseq (n = 3). (**S3B**) IP control of **Figure 4B**. Ponceau S staining of GFP-trap samples showing the corresponding immunoprecipitated products (EGFP = 27 kDa, EGFP-LphD / EGFP-LphD Y392F= 75 kDa, EGFP-RomA=88kDa). (**S3C**) IP control of **Figure 4E**. Ponceau S staining of GFP-trap samples showing the corresponding immunoprecipitated products (EGFP = 27 kDa, EGFP-LphD/ EGFP-LphD Y392F = 75 kDa). (**S3D**) Immunoblots showing the interaction of LphD with KAT7. Co-immunoprecipitation of endogenous KAT7 in HEK293T cells transfected with 3xHA or 3xHA-LphD. Input shows the expression level of endogenous KAT7 and HA-LphD in total lysates, as well as histone H1 (loading control). Co-IP samples were analyzed for the presence of 3xHA-LphD. (**S3E**) IP control of Figure 4F. Ponceau S staining of GFP-trap samples showing the corresponding immunoprecipitated products (EGFP = 27 kDa, EGFP-LphD = 75 kDa and EGFP-RomA=88kDa). (**S3F**) Representative image of western blots for H3K14me, corresponding to Figure 4J. THP-1 cells were pre-treated 18 hours with a KAT7 specific inhibitor (WM-3835) and infected with *L. pneumophila* wild type or *ΔlphD* strain expressing EGFP. Cells were sorted at different times post-infection by FACS, histones were isolated and analyzed by western blot. Histone H1 was used as loading control.

**Supplemental Figure S4: Subcellular localisation of LphD deleted of its putative NLS and H3K18ac and H3K23ac levels in LphD transfected cells.** (**S4A**) Immunofluorescence analysis of subcellular localization of EGFP-LphD full-length and a truncated form for its predicted NLS corresponding to amino acids 2-22 in the N-terminal part of the protein (prediction tool NoD: NucleOlar localization sequence Detector[3]). HeLa cells were either transfected with EGFP-LphD or with EGFP-LphD ΔNLS for 24 hours and then stained with DAPI (cyan), and phalloidin (gray). Scale bars 10 $\mu$m. **(S4B)** Immunofluorescence analysis of H3K14ac levels of cells transfected with EGFP-LphD. HeLa cells were transfected 24 hours with EGFP-LphD and stained for H3K18Ac (red; top) or H3K23Ac (red; bottom), DAPI (cyan), and phalloidin (gray). Scale bars 10 $\mu$m.

Figure S1

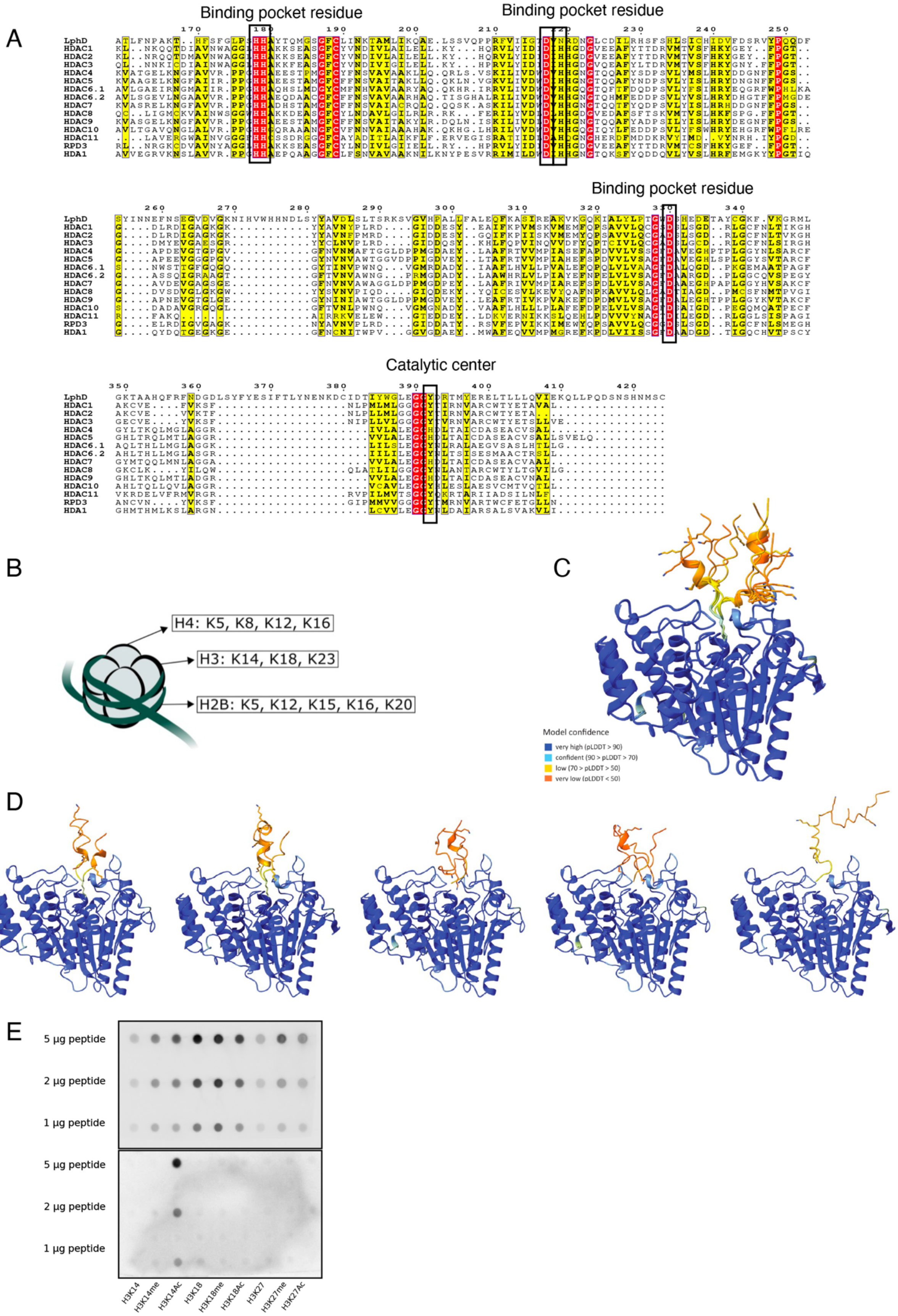
A
Binding pocket residue
Binding pocket residue
Binding pocket residue
Catalytic center
LphD
HDAC1
HDAC2
HDAC3
HDAC4
HDAC5
HDAC6.1
HDAC6.2
HDAC7
HDAC8
HDAC9
HDAC10
HDAC11
RPD3
HDA1
B
H4: K5, K8, K12, K16
H3: K14, K18, K23
H2B: K5, K12, K15, K16, K20
C
Model confidence
very high (pLDDT > 90)
confident (90 > pLDDT > 70)
low (70 > pLDDT > 50)
very low (pLDDT < 50)
D
E
5 µg peptide
2 µg peptide
1 µg peptide
5 µg peptide
2 µg peptide
1 µg peptide
H3K14
H3K14me
H3K14Ac
H3K18
H3K18me
H3K18Ac
H3K27
H3K27me
H3K27Ac

Figure S2

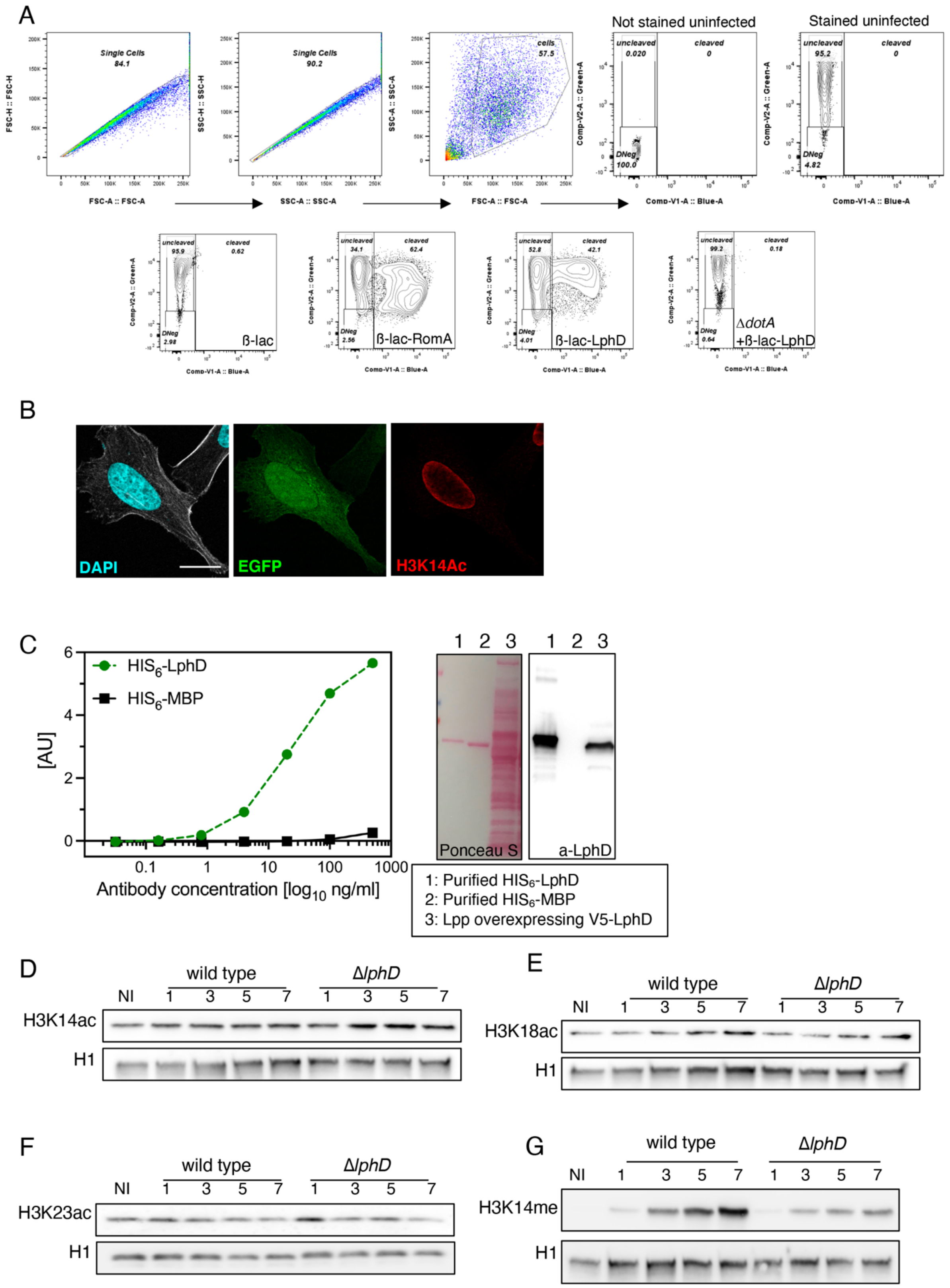

A
Not stained uninfected
Stained uninfected
Single Cells
84.1
Single Cells
90.2
cells
57.5
uncleaved
cleaved
DNeg
FSC-A :: FSC-A
SSC-A :: SSC-A
Comp-V1-A :: Blue-A
ß-lac
ß-lac-RomA
ß-lac-LphD
ΔdotA
+ß-lac-LphD
B
DAPI
EGFP
H3K14Ac
C
HIS6-LphD
HIS6-MBP
[AU]
Antibody concentration [log10 ng/ml]
Ponceau S
a-LphD
1: Purified HIS6-LphD
2: Purified HIS6-MBP
3: Lpp overexpressing V5-LphD
D
wild type
ΔlphD
NI
H3K14ac
H1
E
H3K18ac
F
H3K23ac
G
H3K14me

Figure S3

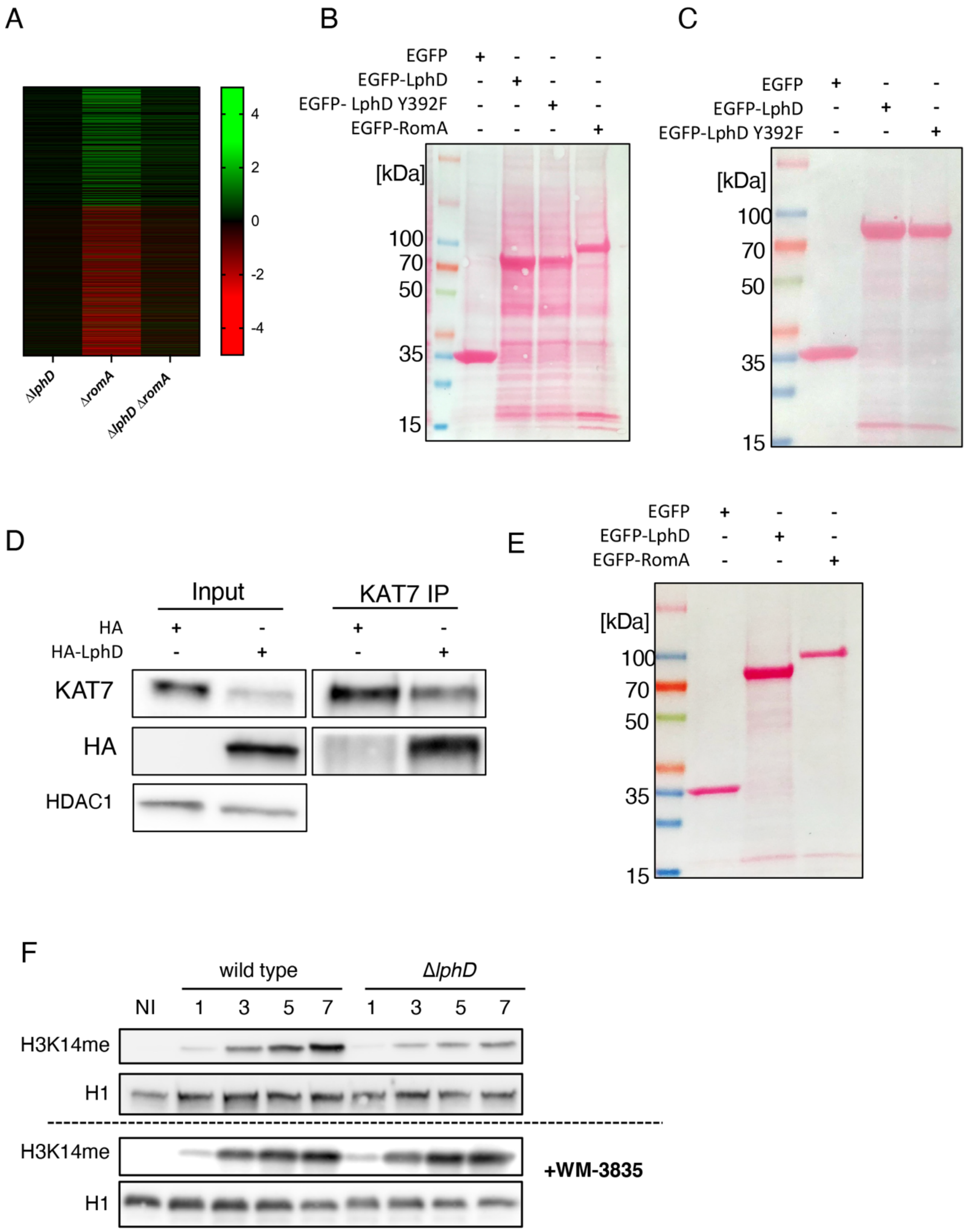
A
4
2
0
-2
-4
ΔlphD
ΔromA
ΔlphD ΔromA
B
EGFP + - - -
EGFP-LphD - + - -
EGFP- LphD Y392F - - + -
EGFP-RomA - - - +
[kDa]
100
70
50
35
15
C
EGFP + - -
EGFP-LphD - + -
EGFP-LphD Y392F - - +
[kDa]
100
70
50
35
15
D
Input
KAT7 IP
HA + - + -
HA-LphD - + - +
KAT7
HA
HDAC1
E
EGFP + - -
EGFP-LphD - + -
EGFP-RomA - - +
[kDa]
100
70
50
35
15
F
wild type
ΔlphD
NI 1 3 5 7 1 3 5 7
H3K14me
H1
H3K14me
H1
+WM-3835

Figure S4

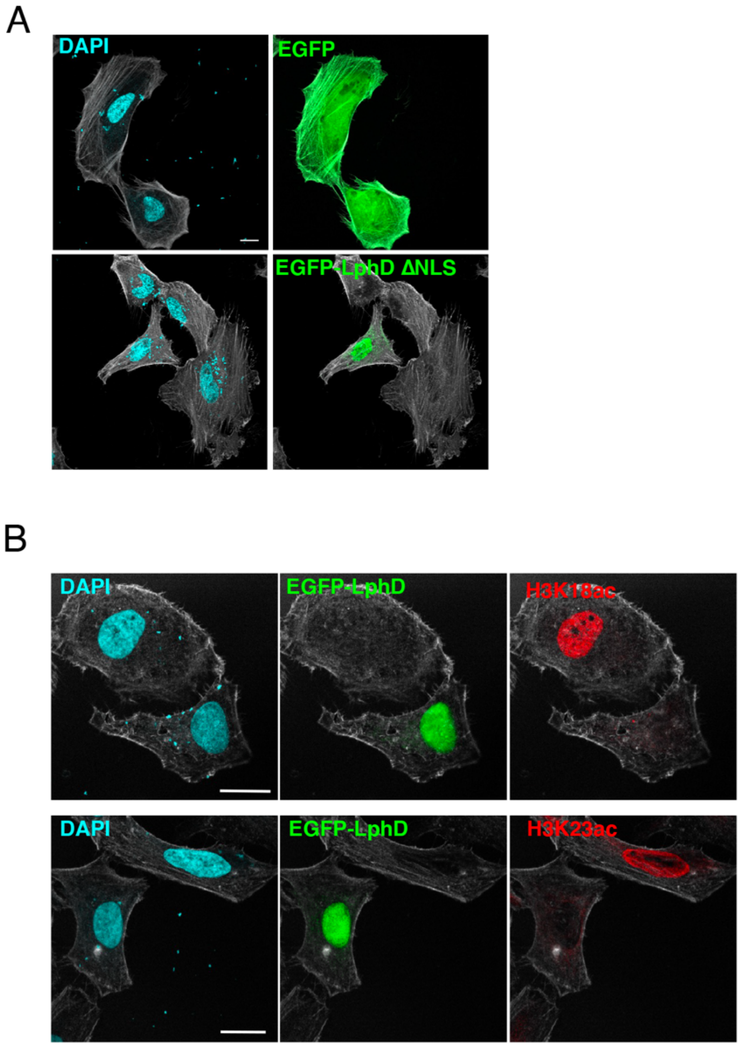
A
DAPI
EGFP
EGFP-LphD ΔNLS
B
DAPI
EGFP-LphD
H3K18ac
DAPI
EGFP-LphD
H3K23ac
